# Deep-Learning Assisted, Single-molecule Imaging analysis (Deep-LASI) of multi-color DNA Origami structures

**DOI:** 10.1101/2023.01.31.526220

**Authors:** Simon Wanninger, Pooyeh Asadiatouei, Johann Bohlen, Clemens-Bässem Salem, Philip Tinnefeld, Evelyn Ploetz, Don C. Lamb

## Abstract

Single-molecule experiments have changed the way we investigate the physical world but data analysis is typically time-consuming and prone to human bias. Here, we present Deep-LASI (Deep-Learning Assisted Single-molecule Imaging analysis), a software package consisting of an ensemble of deep neural networks to rapidly analyze single-, two- and three-color single-molecule data, in particular from single-molecule Förster Resonance Energy Transfer (FRET) experiments. Deep-LASI automatically sorts single molecule traces, determines FRET correction factors and classifies the state transitions of dynamic traces, all in ~20-100 ms per trajectory. We thoroughly benchmarked Deep-LASI using ground truth simulations as well as experimental data analyzed manually by an expert user and compared the results with a conventional Hidden Markov Model analysis. We illustrate the capabilities of the technique using a highly tunable L-shaped DNA origami structure and use Deep-LASI to perform titrations, analyze protein conformational dynamics and demonstrate its versatility for analyzing both total internal reflection fluorescence microscopy and confocal smFRET data.

## INTRODUCTION

Single-molecule spectroscopy has revolutionized how we investigate the mechanism of processes on the nanometer scale. In particular, optical fluorescence imaging allows contact-free investigations of single, dynamic biomolecules, one at a time, in cells, membranes and in solutions. Single-molecule Förster resonance energy transfer (smFRET) in combination with confocal scanning microscopy or total internal reflection fluorescence (TIRF) microscopy probe distances on the nanometer scale (2.5-10 nm). While solution measurements can provide information on sub-millisecond dynamics, measurements with immobilized molecules give access to the temporal evolution of single molecules on the timescale of microseconds to minutes^1^. By removing ensemble averaging, it is possible to directly measure the underlying conformational states and molecular dynamics of biomolecules. Its ability to measure accurate distances and kinetics turned smFRET into a powerful tool for deciphering molecular interaction mechanisms and structures of biomolecules^1–3^. Typically, FRET experiments are performed using two colors and used to probe conformational distributions and distance changes. However, also other single-molecule approaches can be used to investigate small distances changes or interactions (e.g. MIET^4^, GIET^5^, or PIFE^6,7^).

When combining three- or more labels, multi-color FRET can probe molecular interactions between different binding partners and also measure multiple distances simultaneously, i.e., correlated motion within the same molecule^8–10^. However, multi-color analyses remain challenging. Quantitative smFRET data analysis is strongly hampered by experimental restrictions due to (1) low statistics, (2) low signal-to-noise ratio (SNR), or (3) photochemistry. Overcoming these limitations requires large data volumes as very few molecules contain the desired information with suitable quality. Low statistics result from various reasons including molecular events exhibiting slow kinetics or rare transition probability, insufficient labeling efficiency, low SNR, quick photobleaching or spurious background. In addition, arbitrary fluctuations due to unwanted interactions and/or aggregations between binding partners hamper a concise analysis of the underlying state and kinetics.

Various approaches have been developed to overcome these time-consuming burdens, employing user-defined thresholds on the channel count rate, signal-to-noise ratio, FRET values, FRET lifetime, and donor/acceptor correlation.^11–18^ However, setting appropriate thresholds requires a substantial amount of expertise. Depending on the user, the data evaluation is prone to cognitive biases and poses a challenge to reproducible analysis results. Recently, software packages have been published that use deep learning techniques to rapidly automate trace classification and keep user bias to a minimum.^19–21^ In particular, Thomsen et al. comprehensively demonstrated that artificial neural networks can match manual classifications and even outperform conventional methods of commonly used programs to extract valid single-molecule FRET traces.^21^ So far, deep-learning has been solely applied to single-channel and two-color FRET data to categorize the time trajectories for downstream analysis. To study structural dynamics, reflected by changes in intensity and FRET efficiencies, the kinetics are then analyzed separately typically using Hidden Markov Models (HMMs)^22,23^ approaches. Training an HMM requires knowledge of the number of states and modeling of the emission probabilities. Moreover, it assumes that a target state is dependent only on the current state.

While the initial HMM settings are straightforward for simple systems, obtaining the optimal parameters for multi-color FRET becomes a challenging task. To date, only one software package, SMACKS^12^, allows an ensemble HMM for three-color FRET data. As the complexity of the data sets grows, also grows the effort and the required knowledge about the system.

To alleviate the shortcomings of HMM analyses, the hybridization of HMMs with deep neural networks (DNN) have gained popularity.^24–28^ In contrast to HMMs, DNNs are capable of learning higher-order dependencies without prior assumptions about the number and properties of the states. A long-short-term memory (LSTM) neural network was developed to automate stoichiometry determination via photobleaching steps in fluorescence intensity traces.^29^ However, the use of DNNs for extracting quantitative kinetic information from single-molecule data has not yet been explored.

Here, we present the Deep-Learning Assisted, Single-molecule Imaging (Deep-LASI) approach, an ensemble of DNNs with architectures specifically designed to perform a fully automated analysis of single-channel, two-color and three-color single-molecule FRET data. Deep-LASI begins with raw intensity traces and provides corrected FRET efficiencies, state determination, and dwell times without any prior knowledge or assumptions about the system. It classifies each time trace into different categories, identifies which fluorophores are active in each frame to determine FRET correction factors and performs a state transition analysis of the different states in dynamic traces. Deep-LASI also includes optional number-of-state classifiers to estimate the actual number of observed states within one trace. Since the pre-trained neural networks operate locally on each trace, they do not neglect rare events which would be missed in global analysis approaches and lead to inaccurate conclusions. We benchmark the performance of Deep-LASI using ground truth simulations and experimental one-, two- and three-color data using an L-shaped DNA origami structure with tunable dynamic behavior^5,30^. The results are further compared to the manual evaluation of the data and the extracted dwell times obtained with HMM. Finally, we demonstrate the power of Deep-LASI with multiple applications: (1) titration experiments, which would be unfeasible without Deep-LASI; (2) smFRET on a mitochondrial Hsp70 to extract substrate-specific dwell times and conformational states; and (3) the applicability of Deep-LASI to another experimental setup.

## RESULTS AND DISCUSSION

### The Deep-LASI Approach

Deep-LASI utilizes an ensemble of pre-trained deep neural networks designed for the fully automated analysis of one-, two- and three-color single-molecule data including multi-color FRET correction and kinetic analyses. (Figure 1; Supplementary Note 1). The designed input for Deep-LASI is a single-molecule fluorescence intensity trace or traces measured directly using confocal microscopy or extracted from movies using wide-field or TIRF microscopy but is easily expandable to other types of single-molecule datasets. In the case of two-color fluorescence data, continuous wave excitation or ALEX modalities can be analyzed. For three-color smFRET measurements, ALEX data is required. All available channels are fed into a combination of a convolutional neural network (CNN) using the omni-scale feature learning approach and a long short-term memory (LSTM) model (Supplementary Figure SN1.1).

**Figure 1.**
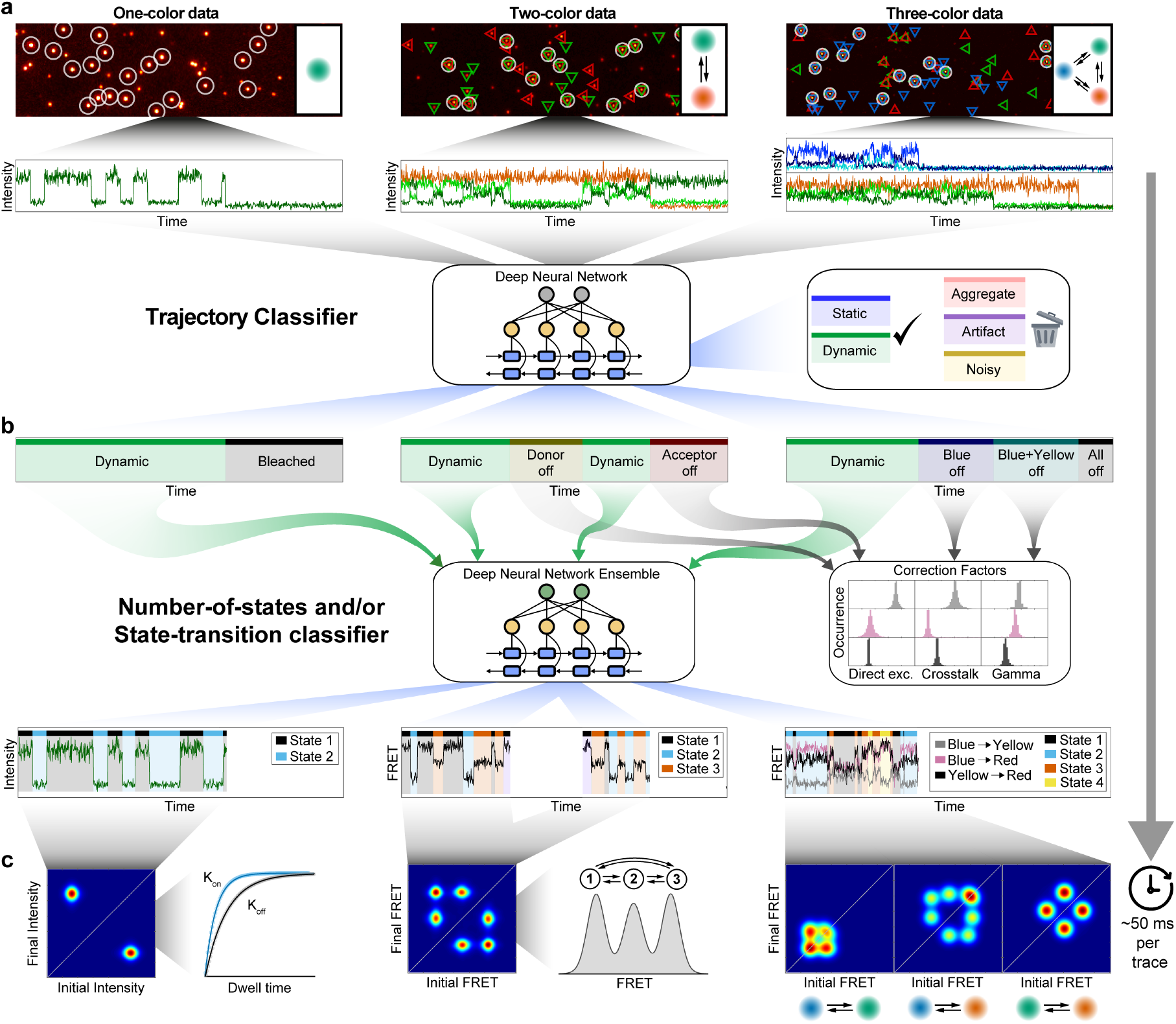
Overview of data extraction, evaluation, and analysis using Deep-LASI. **(a)** Single-molecule data of up to 3 separate channels after direct and alternating laser excitation are identified, extracted, and presorted for further analysis. Each frame within the time traces is classified into categories using a hybrid CNN-LSTM. **(b)** A second hybrid CNN-LSTM evaluates the kinetics and state information in the presorted data. The photobleaching information can be used for determining the correction factors framewise or state-wise to obtain accurate FRET values between 2 and 3 fluorophores. **(c)** Next, the interconversion rates between underlying states and absolute, distance-related FRET values are extracted from multi-color data sets.

Deep-LASI extracts spatial and temporal sequence features simultaneously and classifies every frame into a specific category (Figure 1a). Building upon Deep-FRET for two-color FRET analysis^21^, we separate the traces into a ‘dynamic’, ‘static’, ‘noisy’, ‘artifact’, ‘aggregate’, or photobleaching category (see Supplementary Note 2 for details). The total number of categories depends on the number of input channels, i.e., the number of dyes (and alternating light sources) used in the experiment. Traces containing random artifacts, aggregates, or high noise are excluded from further analysis. The final output of the state classifier provides an estimation of the probability for each category. The summed probabilities over all unbleached frames serve as confidence levels for each trace. Here, user-defined thresholds can be set to increase or decrease the tolerance towards non-ideal traces to be included in further analysis. In contrast to previous networks, Deep-LASI detects photobleaching events of individual dyes and, therefore, allows the calculation of correction factors obtainable for that molecule. Traces showing no apparent state transition are classified as static and can be included, e.g., in the final corrected FRET histograms.

All sections in each trajectory identified as dynamic are transferred to the state classifier network (Figure 1b), which is designed to detect transitions based only on the intensity data and not via the FRET efficiency. The state classifier assigns every frame to one of the multiple states present in a dynamic trace section and again provides a confidence value of state occupancy that can be used for additional thresholding. Given the state transition classifications, a transition density plot (TDP) is calculated and the kinetic rates of all identified states can be extracted by fitting the corresponding dwell time distributions (Figure 1c). Starting from trace extraction, the TDP marks the first necessary point of human intervention, i.e., the manual selection of state transitions and the fitting procedure. Thus, user bias is kept to a minimum. No assumptions are needed regarding the number of states, state-specific emission probabilities, or other settings required for conventional methods such as Hidden Markov Models (HMM). Of course, as for any deep learning algorithm, the output of the analysis is dependent on the quality and appropriateness of the training data used. Depending on (1) the total number of frames, (2) the yield of valid frames, (3) the computer performance, and (4) the desired confidence threshold, a given data set can be fully categorized on a time scale of 20-100 ms per trace.

### Training of Deep-LASI

To use Deep-LASI for analyzing single molecule data, we first trained the trace-classifier network on appropriate data sets. As the noise sources in single-molecule fluorescence intensity data are well understood, simulated traces are well suited for training the neural network. In addition, it has the advantages of being able to minimize biases and quickly retrain neural network models for adjusting to specific circumstances. The training data sets were designed to cover a wide range of experimental conditions and all possible FRET combinations. A detailed description of the program architecture, simulations, training data sets and benchmarking can be found in the *Materials and Methods* section as well as in Supplementary Notes 1 - 4.

Deep-LASI contains a total of 16 pre-trained deep neural networks for state classification. Four models account for the classification and segmentation of time trajectories obtained from measurements using single-channel data acquisition, two-color FRET with continuous wave excitation, two-color FRET with ALEX, and three-color FRET with ALEX. For each type of experiment, we provide three state-transition-classifiers trained on either two, three or four observed states, which take the output category ‘dynamic’ as the input. Note, acceptor intensity after direct excitation does not contain relevant kinetic information and is not used in the state classifier networks. In addition, a deep neural network is provided that has been optimized for detecting the actual number of observed states and can be utilized for model selection. These neural networks are not essential in the automated analysis process but can serve as a safeguard against trajectories that may be out of the scope of the state transition classifiers.

### Performance of Deep-LASI

A common approach to benchmark classifier models is using ground truth labeled data and calculating confusion matrices, which summarize the correct and incorrect predictions. For every trained model (using ~ 200,000 traces), we generated approximately 20,000 new traces for testing, which were not part of the training data set. Each of the validation data sets was then fed into the corresponding model. The output predictions were compared to the ground truth labels for every frame to obtain the percentage values of true positive, false positive and false negative classifications. All trace classifier models achieve a minimum combined precision of 97 % in predicting smFRET categories, i.e., ‘static’ or ‘dynamic’, and 96 % in predicting non-smFRET categories (Supplementary Figure SN3.1 and SN3.2).

Our number-of-states and state-transition classifiers were benchmarked analogously. For the number-of-state classifiers, two states can be distinguished from multi-state trajectories with at least 98 % precision whereas four states are predicted with the lowest precision of 86 % for the single-channel model (Supplementary Figure SN3.3). For the state-transition classifiers, the states can be identified with accuracies of ≥98%, ≥90% and ≥78% for two-state, three-state and four-state models respectively (Supplementary Figure SN3.4). The comparison between all state-transition classifiers reveals a clear trend of decreasing accuracies with an increasing number of states and increasing accuracy with an increasing number of available channels. This is expected since a higher number of states have a larger probability of lower contrast, and a higher number of channels improves the robustness towards uncorrelated noise. Since confusion matrices do not reveal any underlying dependencies, we additionally benchmarked the state transition classifiers with HMM by calculating the accuracy of dwell time recovery and the precision of the state label prediction for a broad range of noise levels, FRET state differences and dynamic time scales (Supplementary Figure SN3.5). Overall, the performance of state classifiers is at least on par with HMM at low noise levels and outperforms HMM at high noise levels by up to 30 %.

### Deep-LASI analyses of DNA origami structures

Next, we benchmarked the potential of Deep-LASI to automatically analyze experimental data obtained from DNA origami structures. DNA origami is extensively used in bionanotechnology and has the advantage of being programmable with high precision and controllability. In particular, we choose an L-shaped DNA nanostructure with a dynamic, fluorescently labeled 19 nucleotide (nt) single-stranded DNA pointer^5,30^. FRET efficiencies and kinetic rates could be tuned by varying the position and complementary sequence length of binding strands on the DNA origami platform. We designed various DNA origami structures with one-, two-, and three-color labels and measured them on the single-molecule level.

#### Single-Color Experiments

In the first assay, we probed one-color single-molecule kinetics where the flexible pointer was labeled with Cy3B at the 3’-end. Two complementary binding sites with 8 nt complementary nucleotides containing a 1 nt mismatch at the 5’-end (referred to as 7.5 nt) were placed about 6 nm below and above the pointer position (Figure 2a). Binding occurred by spontaneous base-pairing to single-stranded protruding strands. A single red dye, Atto647N, acting as a quencher, was attached about 3 nm aside from the upper binding site (state 1). Figure 2b shows an exemplary intensity trajectory of Cy3B classified as dynamic until photobleaching was detected by the trace-classifier with two corresponding states determined by the state-classifier as the linker moves up and down.

**Figure 2.**
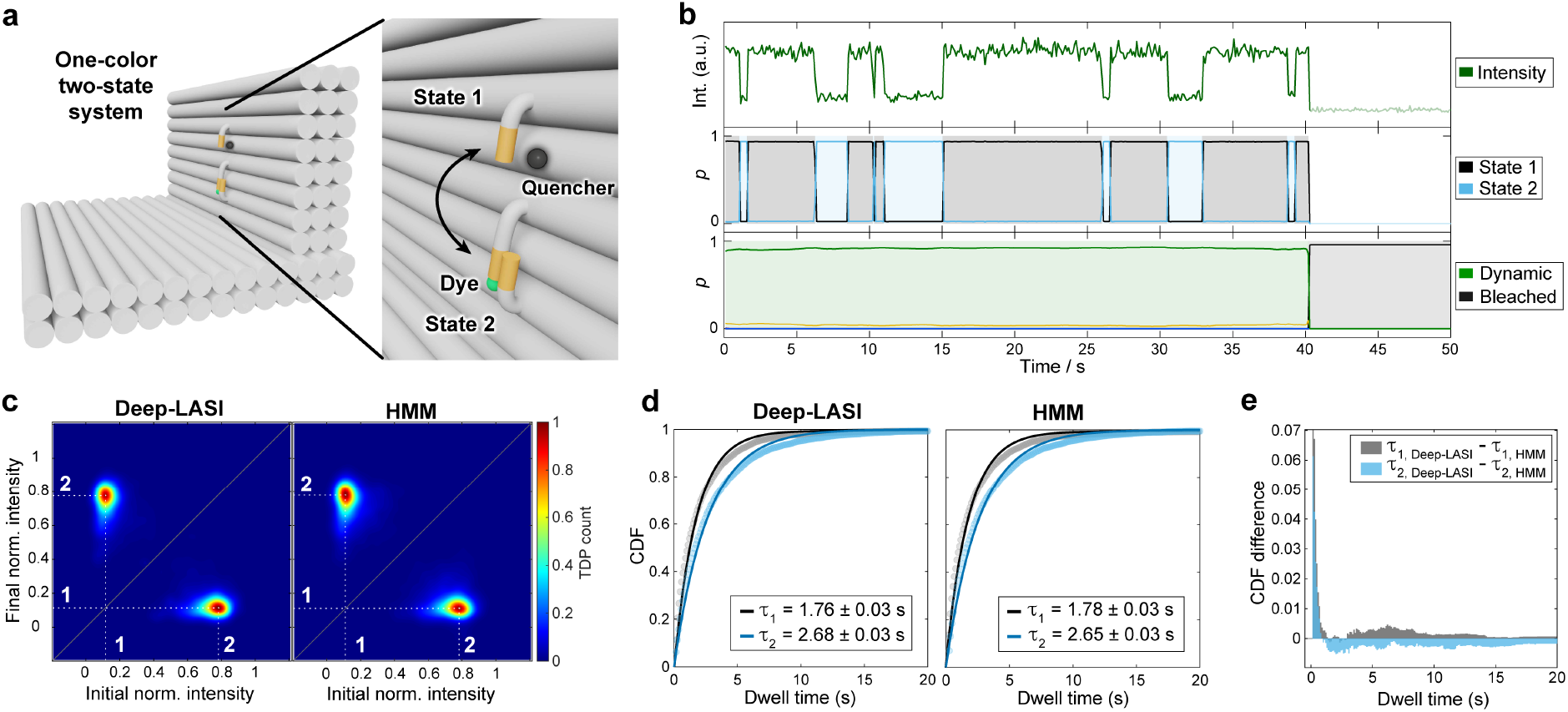
State analysis of single-color single-molecule data. **(a)** Sketch of the used L-shaped DNA origami structure with a single fluorophore (Cy3B) attached to a flexible tether. The zoom-in shows the two single-stranded binding sites (orange) in close and distant proximity to a quencher dye (Atto647N) bound to the DNA origami structure. **(b)** Representative time transient for a DNA origami structure with 7.5 nt binding strands after classification and kinetic evaluation by Deep-LASI. **(c)** Transition-density plots depicting the interconversion events between the two detected states 1 and 2 after trace kinetics evaluation by Deep-LASI (left) and using a Hidden-Markov Modeling (HMM) analysis (right). Both approaches identify identical states. **(d)** The mono-exponential fits obtained by both methods reveal equivalent dwell times of approximately 1.75 and 2.65 seconds for the upper (State 1) and lower (State 2) binding sites respectively. **(e)** A comparison of the cumulative dwell-time distribution was determined using Deep-LASI and HMM. Deep-LASI is already sensitive at time scales on the order of the acquisition time. The average difference is less than 1% between both methods.

We compared the results from Deep-LASI with a Hidden-Markov-Model analysis (HMM) trained on the same dataset. Since the state classifier does not directly predict a pre-trained intensity value for each state, the TDP was generated by averaging the normalized intensity between transitions. Both methods yield identical TDPs (Figure 2c). The residence time of the DNA tether in both states was determined by fitting the cumulative dwell-time distribution functions (CDFs) derived from the state-classifier of Deep-LASI with a mono-exponential fit and compared it with the results of HMM. The dwell times of 1.76 s versus 1.78 s (State 1) and 2.68 s versus 2.65 s (State 2) for Deep-LASI and HMM, respectively, are in excellent agreement (Figure 2d). The differences between the CDFs obtained by Deep-LASI and HMM (Figure 2e) indicate that Deep-LASI can identify even fast transitions close to the frame time in contrast to HMM. The overall difference at longer dwell times remains well below 1%, which proves that Deep-LASI obtains identical results to HMM with negligible differences in the extracted rates. Interestingly, although the DNA binding strands are identical in sequence and length, there are clear differences in the dwell times. We attribute this to an inherent bias in the equilibrium position of the DNA pointer.

#### Dual-Color Experiments

Next, we investigated Deep-LASI’s capabilities to study two-color FRET assays with two states and compared the results with a pure manual evaluation of the same data. Here, both donor and acceptor signals from the same DNA origami sample system as shown in Figure 2a were analyzed (Figure 3a). TIRF measurements were performed using msALEX^31^ yielding donor signal (Cy3B, Channel D_ex_D_em_), sensitized emission (Channel D_ex_A_em_) and acceptor signal (Atto647N, Channel A_ex_A_em_) to obtain information about acceptor photobleaching and direct excitation. Figure 3b shows a fully classified example trace with the signals on top and the derived FRET trace below. From the trace classifier, Deep-LASI identified dynamic sections and individual photobleaching events (Figure 3b; bottom). The dynamic section was further classified in the state-transition classifier according to their state occupancy using only the two channels of the donor and acceptor intensity after donor excitation (Figure 3b; middle). The channel of acceptor excitation and detection does not serve as input for the state transition classifier since it does not contain valuable kinetic information. From a total of 6100 recorded traces in the data set, 1499 traces were classified as dynamic smFRET trajectories with at least one transition.

**Figure 3.**
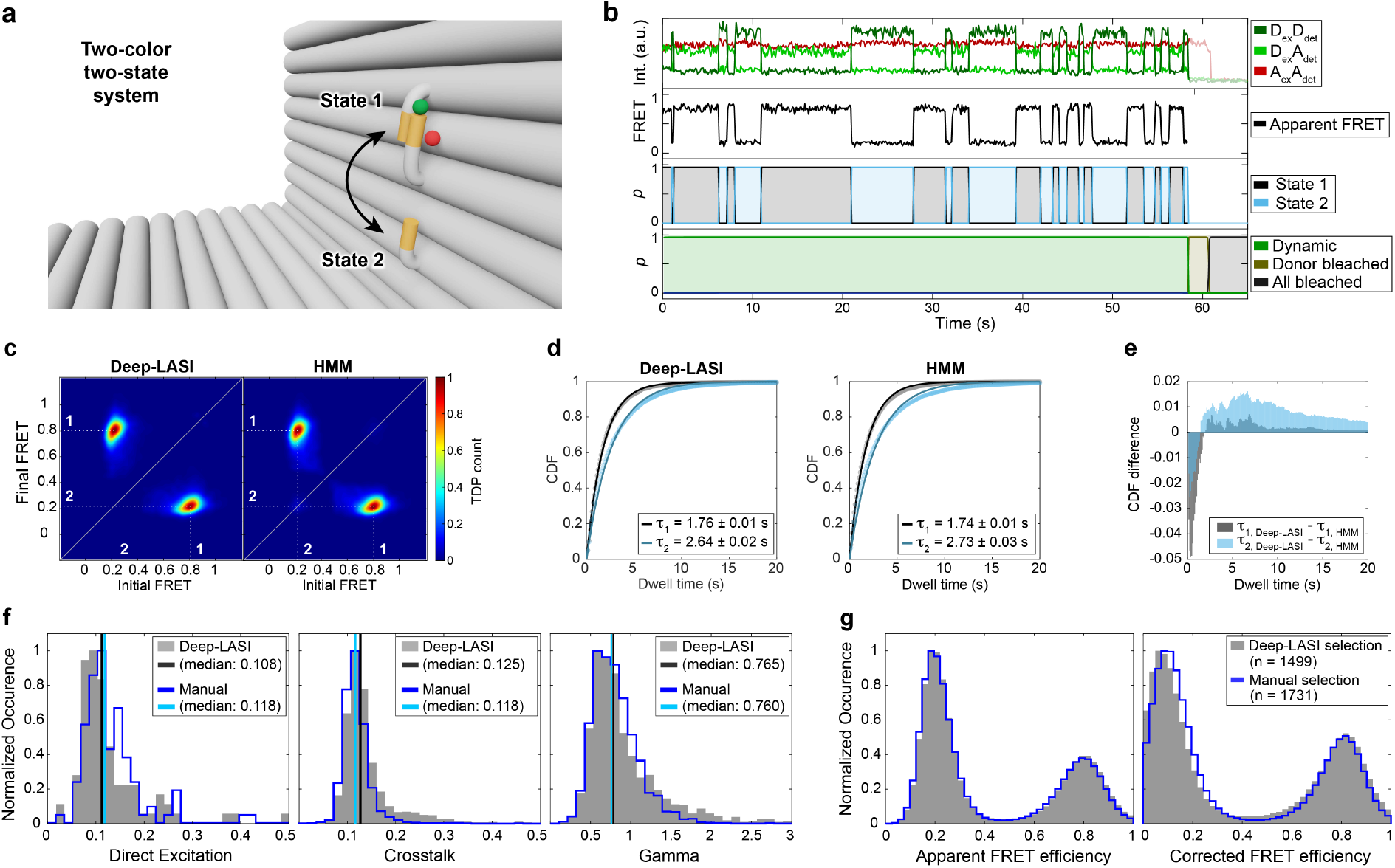
Single-molecule analysis of two-color FRET data. Experiments were performed with DNA origami structures exhibiting two (a-e) and three (f-j) FRET states. **(a)** Zoom-in of an L-shaped DNA origami structure labeled with Atto647N and Cy3B. The donor is attached to the flexible tether with a 7.5 nt overhang between the pointer and two single-stranded binding sites. FRET is expected between a high FRET State 1 (12 o’clock) and a low FRET State 2 (6 o’clock). **(b)** Representative single-molecule and apparent FRET trace after alternating red-yellow (RY) laser excitation. Deep-LASI classifies the trace and determines the underlying state for each frame. **(c)** TDPs determined using Deep-LASI (left) and HMM (right) are shown revealing two interconverting states with apparent FRET values of 0.8 and 0.2. The two states are labeled in white. **(d)** CDFs extracted from the TDPs shown in panel (c) yielding dwell times of 1.76 s and 2.64 s, respectively. Total amount of transitions: 15958. **(e)** A comparison of the cumulative dwell-time distribution determined using Deep-LASI - HMM for *τ*1 (gray) and *τ*2 (cyan). **(f)** Histograms of trace-wise determined correction factors for direct excitation, crosstalk and detection efficiency, either derived automatically by Deep-LASI (gray histograms, median in black) or determined manually (blue lines, median in cyan) (see Supplementary Note 5). **(g)** Apparent (left) and corrected (right) frame-wise smFRET efficiency histograms for 1499 dynamic traces out of a total of 6100 traces. The states have corrected peak FRET efficiencies of 0.07 and 0.81. The histograms from traces selected by Deep-LASI are shown in gray and by manual selection in blue.

The same traces were also sorted manually and the 1731 selected dynamic traces were analyzed using HMM^32^ (see Supplementary Note 5 for details). TDPs from the state-transition classifier and from the HMM analysis are nearly identical (Figure 3c). Also, the corresponding dwell times, determined via mono-exponential fits to the CDFs, are similar (Figure 3d) and correspond to the expected dwell times of the one-color sample shown in Figure 2 (~1.75 s for state 1 and 2.68 s for State 2). Again, a comparison of the CDFs from Deep-LASI and HMM indicates that Deep-LASI is better at recognizing fast transitions (Figure 3e).

To determine the distance between both dyes in the two FRET states, the smFRET data needs to be corrected. Deep-LASI uses the frames classified as photobleached to automatically derive the correction factors necessary for an accurate FRET calculation^1,33,34^. In the manual analysis, the relevant regions are selected by hand (Figure 3f, Supplementary Note 5). The correction factors agree within ~3 %. Using the derived correction factors, the correct FRET efficiency is determined. The apparent (left) and corrected FRET histograms (right) of the Deep-LASI (gray histograms) and manually (blue lines) selected traces are shown in Figure 3g. There is excellent agreement between the Deep-LASI and manually analyzed apparent FRET histograms. The difference between the corrected histograms is due to the difference in the correction factors determined and applied from the two analyses. In this case, as Deep-LASI classifies photobleaching on a per-frame basis, more frames can be used for determining the correction factors and is thus, most likely, more accurate here. The corrected peak FRET efficiencies are 0.81 and 0.82 (State 1) and 0.08 and 0.14 (State 2) for Deep-LASI and manual evaluation, respectively, and correspond to distances of 53 and 53 Å, and 103 and 92 Å (assuming an *R*_0_ of 68 Å^7^).

#### Three-color experiments

We then tested the performance of Deep-LASI for analyzing three-color data by labeling the DNA origami structure with an additional blue dye, Atto488, at ~3 Å distance to the binding site for State 2 (Figure 4a). The labeling sites of the yellow (Cy3b) and red (Atto647N) dyes were left unchanged to provide consistency with the previous two-color experiments. The use of three FRET pairs provides three distances simultaneously and allows the resolution of states that are degenerate for two-color FRET.

**Figure 4.**
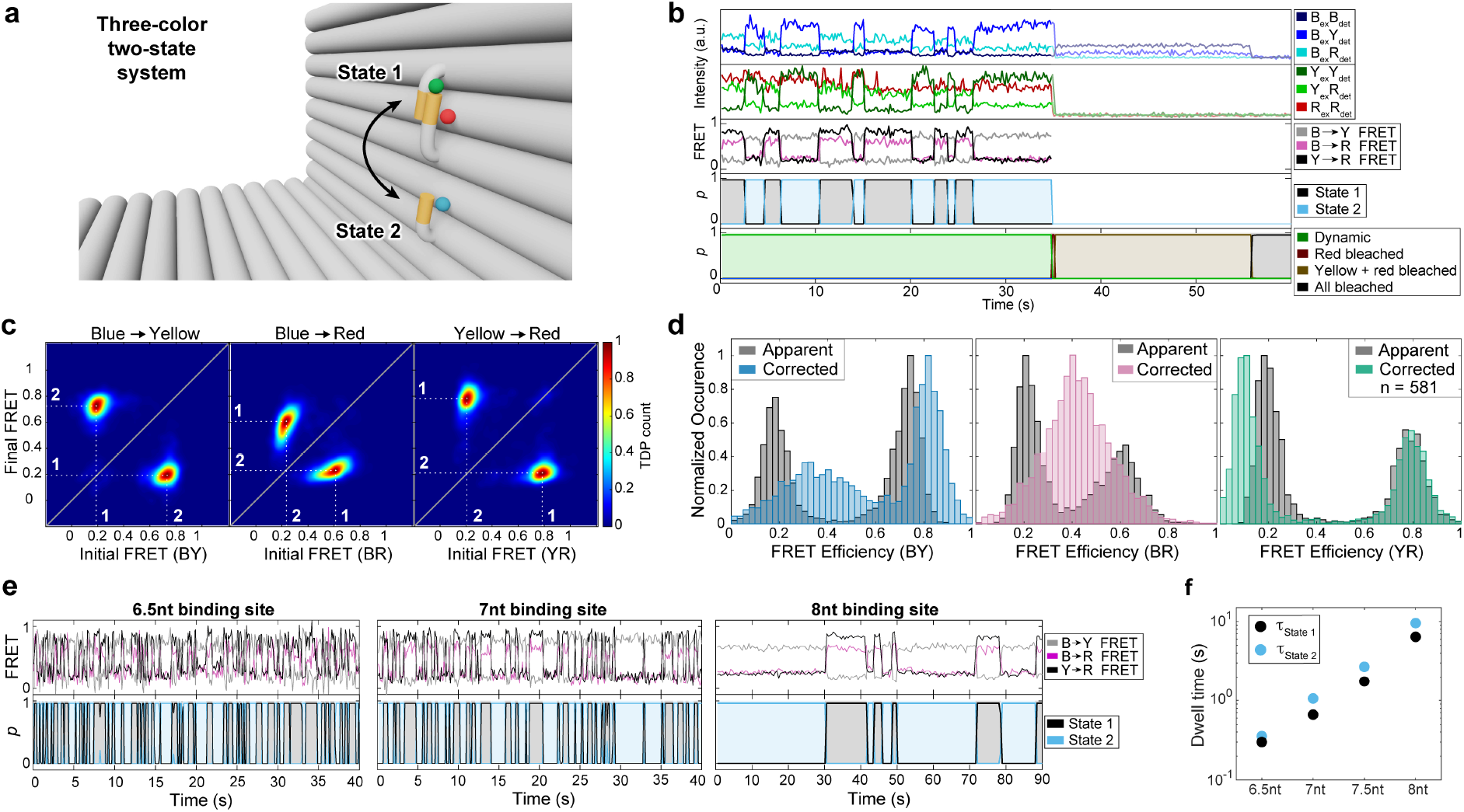
Single-molecule analysis of three-color FRET data. Experiments were performed with DNA origami structures exhibiting two FRET states. **(a)** Zoom-in of an L-shaped DNA origami structure labeled with Atto647N, Cy3B and Atto488. Cy3B is attached to the flexible tether. Atto647N and Atto488 are bound on the DNA origami structure near the binding site at the top (12 o’clock; State 1) and bottom (6 o’clock; State 2), respectively. **(b)** Representative single molecule intensity and FRET trajectories for binding sites with 7.5 nt overhang after alternating red-yellow-blue laser excitation. The upper panel shows the intensity in the blue, yellow and red channels after blue excitation. The second panel shows the intensity in the yellow and red channels after yellow excitation and the red intensity after red excitation. The middle panel shows the corresponding FRET efficiencies for the three dye pairs. The fourth and fifth panels show the output of the Deep-LASI analysis for state-transition and trace classification respectively. **(c)** TDPs of the apparent FRET efficiency states are shown. One sees an apparent distance change for all three FRET pairs (BY (left), BR (middle), and YR channel (right)) with dwell times of 1.75 s and 2.69 s seconds for the upper and lower binding site, nearly identical to the two-color DNA origami structures (Figure 3c). **(d)** Frame-wise weighted state-wise apparent (gray) and corrected (color) smFRET efficiency histograms of the three FRET pairs: BY (left), BR (middle), and YR channel (right). As expected from the design, the accurate FRET efficiency between the blue and red dye is static and amounts to 0.36. As the position of Cy3B changes from State 1 to State 2, the accurate FRET efficiency changes from 0.36 to 0.81 in the BY channel and from 0.81 to 0.08 in the YR channel. **(e)** *Upper panel:* Representative three-color smFRET traces for binding sites with 6.5 nt (7 nt with 1 nt mismatch), 7 nt and 8 nt overhangs after alternating RYB laser excitation. *Bottom Panel*: The corresponding state determined by Deep-LASI. **(f)** Extracted dwell times of the lower (blue) and upper position (red) for 6.5 nt (τ_2_: 0.31 s, τ_2_: 0.4 s), 7 nt (τ_1_: 0.66 s, τ_2_: 1.05 s), 7.5 nt (τ_1_: 1.75 s, τ_2_: 2.69 s) and 8 nt overhangs (τ_1_: 6.41 s, τ_2_: 9.54 s) (see Supplementary Figure SN6.1 for more details).

Using the six available intensity traces, each frame is categorized by the fluorophores that are active and whether the trace is static, dynamic or should be discarded. As the acceptor intensity after acceptor excitation (R_ex_R_em_) does not contain valuable kinetic information, the other 5 intensity channels for dynamic traces (before photobleaching) are given as input for the state transition classifier (Figure 4b). Movement of the flexible tether results in an anti-correlated change in the FRET efficiency of blue to yellow (BY) and yellow to red (YR), visible in the apparent FRET panel of the example trace in Figure 4b. For each FRET pair, a TDP can be calculated, which allows the assignment of the state number to the actual FRET populations (Figure 4c). Note, the apparent FRET efficiency of blue to red (BR) varies with the YR FRET efficiency due to the different energy transfer pathways taken upon blue excitation. Deep-LASI classifies a state regardless of which dye is undergoing a transition, i.e., the extracted dwell time distribution of a given state is the same for all FRET pairs when there is no overlap of multiple states in the TDP. The dwell times for states 1 and 2 match with those for the one-color and two-color samples, which indicates that the transition rates are not influenced by the acceptor dyes close by (Figure 2d, Figure 3d, Supplementary Figure SN6.1). From a total of 2545 recorded molecules, 581 were classified as valid, dynamic three-color FRET traces. The uncorrected, framewise smFRET histograms of BY, BR and YR FRET pairs are very similar to those from the 694 manually selected traces (Supplementary Figure SN4.1a).

As for two-color FRET, Deep-LASI automatically determines all correction factors obtainable per trace depending on which dyes are photoactive. The results of the automated extraction of correction factors are summarized and compared to manually derived correction factors in Supplementary Figure SN4.1b. The corresponding apparent und state-wise, corrected FRET efficiency histograms for each FRET pair are shown in Figure 4d. While the YR FRET efficiency can be directly calculated, the corrected BY and BR FRET efficiencies are subjected to higher uncertainties due to the large number of correction factors involved (see Supplementary Note 5). In particular, their dependency on the YR FRET efficiency leads to the broadening of the distributions. To minimize this influence, we perform the correction using the state-averaged FRET efficiencies to avoid the use of outliers during the correction. After correction, the FRET efficiencies of State 1 (0.81) and State 2 (0.08) for the YR FRET pair are virtually identical as for the two-color system. For the BY FRET pair, State 1 and State 2 correspond to peak FRET efficiencies of 0.36 and 0.81, respectively. As expected, the two populations of the apparent BR FRET efficiency merge into one static population in the corrected histogram with a peak FRET efficiency of 0.36.

To probe the performance of the kinetic analysis from Deep-LASI, we used the tunability of the L-shaped DNA origami structure to vary the timescale of the dynamics. In addition to the 7.5 nt binding sites (Figure 4a-d), we measured three samples using binding sites of length 7 nt with a 1 nt mismatch (referred to as 6.5 nt), 7 nt, and 8 nt (Figure 4e). The summary of all extracted dwell times (Figure 4f, Supplementary Figure SN6.1) shows an exponential increase in the dwell times of both states with increasing binding site lengths ranging from 0.1 s to 3.3 s. Considering the camera exposure times of 50 ms (6.5 nt, 7 nt and 7.5 nt data sets) and 150 ms (8 nt data set), a dwell-time to frame-time ratio ranges from 10 (6.5 nt State 1) to 62 (8 nt, State 2).

To test Deep-LASI with more complex dynamics with multiple states, we constructed a three-state system with three-color labels using 7 nt binding strands at positions 6 and 12 o’clock and an additional 7.5 nt complementary binding strand at 9 o’clock (Figure 5a). An example trace containing all possible transitions identified by Deep-LASI is shown in Figure 5b. The TDP of the BY FRET pair (Figure 5c, left panel) yields clearly distinguishable populations, while the TDP of the YR FRET pair (Figure 5c, right panel) shows a degeneracy of state 3 transitions. Using the BY TDP, we determined the dwell time distributions with residence times between 0.65 and 1.43 s (Supplementary Figure SN6.2). The three states are well-resolved in the framewise apparent BY FRET histogram, while state 2 and state 3 are degenerate for the BR and YR FRET pairs (Figure 5c). Applying all correction factors yields peak YR FRET efficiencies of 0.81 (state 1), 0.08 (state 2) and 0.19 (state 3). Upon correction, States 1 and 3 in the BY FRET histogram merge into a broad degenerate FRET population. However, using the state information for all three fluorophores allows us to separate out the BY FRET histograms of the individual states.

**Figure 5.**
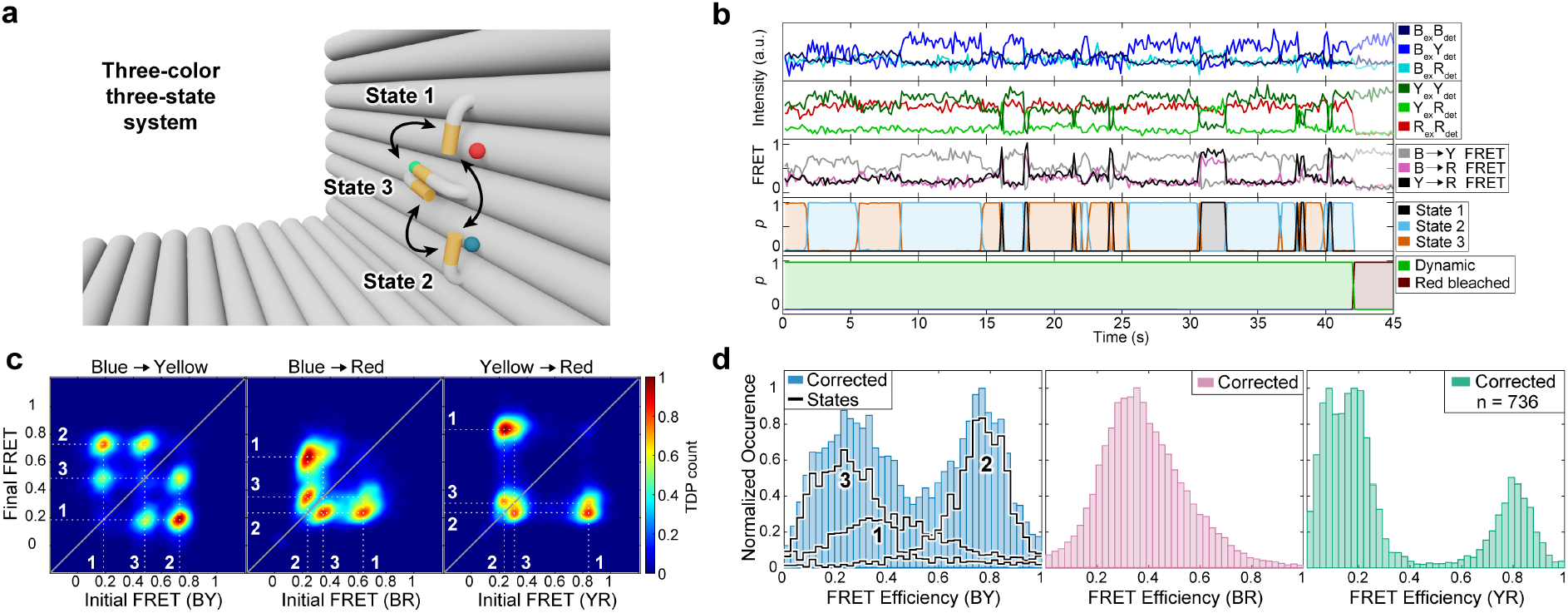
Single-molecule analysis of three-state, three-color FRET data. **(a)** Zoom-in of the L-shaped DNA origami structure with an additional binding site for the tether (State 3 at 9 o’clock). **(b)** A representative single-molecule three-color FRET trace and apparent FRET for the 3-state system. The upper panel shows the intensity in the blue, yellow and red channels after blue excitation. The second panel shows the intensity in the yellow and red channels after yellow excitation and the red intensity after red excitation. The middle panel shows the corresponding FRET efficiencies for the three dye pairs. The fourth and fifth panels show the output of the Deep-LASI analysis for state-transition and trace classification respectively. **(c)** Transition density plots of the apparent FRET efficiency states are shown for each FRET pair revealing an interconversion between 3 binding sites. **(d)** Frame-wise weighted state-wise corrected smFRET efficiency histograms. Corrected, distance-related FRET values are best resolved for the YR pair (cf. Figure 3i-j), showing three populations at 0.81, 0.19 and 0.09. The BY FRET shows one population at 0.8 corresponding to state 2 and a broad population at 0.3 of the states 2 and 3. Individually corrected states are indicated with the highlighted lines, showing the actual BY FRET efficiencies of state 2 (0.4) and state 3 (0.21). The apparent FRET states for the BR channel merge into one broad, static state with a value of 0.35.

For three-color FRET, the corrected BY and BR FRET efficiencies depend on the YR FRET efficiency and the additional corrections broaden the population. However, even though the data may be noisier, three-color experiments contain additional information, which typically allows one to resolve states that are degenerate in two-color experiments. This is exemplified in two-color FRET experiments on the same construct missing the blue fluorophore near the 6 o’clock binding site (Supplementary Note 6.3). For distinguishable states, the determined corrected FRET efficiencies and kinetic rates from two- and three-color experiments are the same. However, three-color FRET experiments enable the lifting of this degeneracy between states 2 and 3. To minimize the influence of the increased noise in three-color experiments, it is advantageous to analyze the data in proximity ratio and only convert it to corrected FRET efficiencies when necessary.^9^ Deep-LASI can rapidly classify a large number of molecules and quickly provide an overview of multi-state dynamics with easy access to the kinetic information.

Lastly, we compared the performance of Deep-LASI with other kinetic analysis routines that have been recently published in a multi-laboratory study. We chose to analyze the two-state datasets as these require no user input and the analysis can be performed without bias. Deep-LASI returned values corresponding to the ground truth for the simulated dataset and close to the average values obtained for the experimental dataset (Supplementary Figure SN6.4).

### Further applications of Deep-LASI

After extensively benchmarking Deep-LASI, we applied Deep-LASI to various single-molecule data sets originating from biophysical assays, protein samples and experimental systems beyond TIRF microscopy. With the speedup in analysis time from days to minutes, experiments become possible that would have been unthinkable when performing the analysis manually. One example is a titration experiment where the biochemical conditions are varied. Here, we measured the influence of glycerol concentration on the dynamics of the 3-colored L-shaped DNA origami introduced in Figure 4a with 7.5 nt overhangs. Interestingly, we observed a decrease in residence time in both states with increasing glycerol concentrations (Figure 6a,b). Dwell times start at 1.75 s (state 1) and 2.69 s (state 2) for pure imaging buffer and decrease down to 0.62 s and 0.85 s in buffer containing 30 % (v/v) glycerol. The multi-fold increase in binding kinetics can be explained by a reported destabilization of base-pairing due to changes in the ssDNA hydration shell^35^ and concomitantly disturbed hydrogen bonding due to the osmolyte-DNA interaction. The melting enthalpy and melting temperature decreases linearly with glycerol concentration at about 0.2 °C per % (v/v)^36,37^ in line with our observations (Figure 5b). With Deep-LASI at hand, local screening and time-consuming parameterization of imaging conditions becomes feasible.

**Figure 6.**
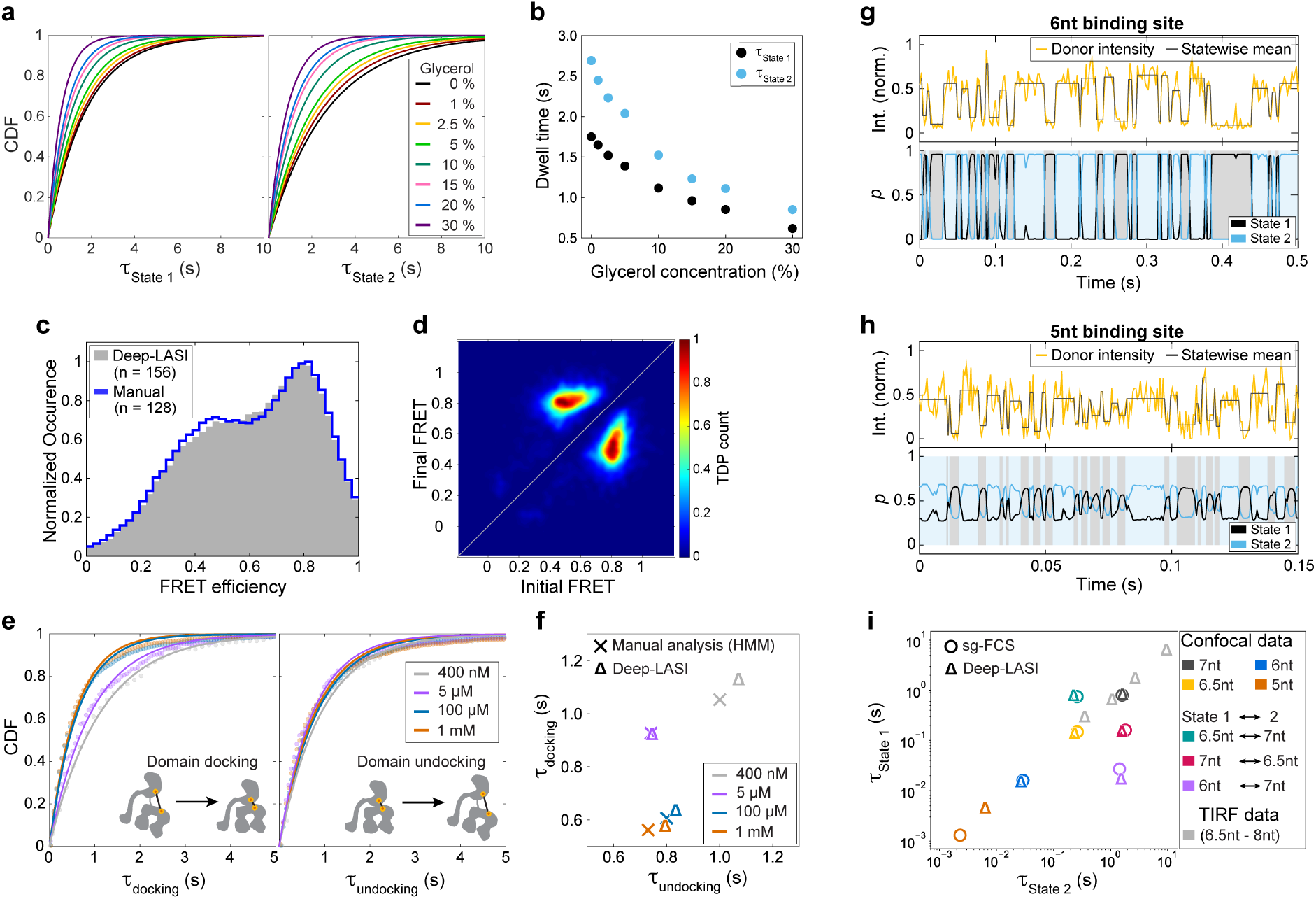
Use of Deep-LASI on titration experiments, protein data, and confocal data. **(a-b)** *3cFRET: Tuning the dissociation thermodynamics between protruding ssDNA strands by osmolytes.* **(a)** The cumulated dwell times, determined by mono-exponential fits, of state 1 (left) and state 2 (right) for the L-shaped DNA origami structure introduced in Figure 4a decrease with increasing glycerol concentration. **(b)** A plot of the dwell times for both states as a function of glycerol concentration is shown. **(c-f)** *2cFRET: Probing domain-domain interactions in Ssc1, a mitochondrial Hsp70.* **(c)** Frame-wise smFRET distributions of Hsp70 molecules in the presence of 1 mM ADP classified as dynamic by Deep-LASI (gray) and evaluated manually (blue) from a total of 5035 traces. **(d)** The TDP generated by Deep-LASI corresponding to the data plotted in panel (c) showing the interconversion between the undocked (~0.5) and docked (~0.8) conformation. **(e)** The cumulative probability distribution function and mono-exponential fits to the dwell time distributions derived by Deep-HMM for domain docking (left panel) and domain undocking (right panel) as a function of ADP concentration. **(f)** A comparison of the average dwell times extracted by Deep-LASI (triangles) and manually evaluated (crosses) using HMM. Deep-LASI yields similar dwell time values and the same trend as previously published^38^. *(g-i) 1cFRET: Deep-LASI analysis of ssDNA binding kinetics measured using confocal microscopy.* **(g)** A confocal trace (with 2 ms binning) of the DNA origami structure introduced in Figure 2a with 6 nt binding sites and corresponding state labels predicted with high confidence. **(h)** A confocal trace (with 0.6 ms binning) of a DNA origami structure with 5 nt binding sites and predicted state labels. Due to the low SNR of the data, the confidence output of Deep-LASI is low and reaches its limit. **(i)** An overview of the mean dwell times obtained from confocal data for various binding site lengths using sg-FCS^39^ (circles) and Deep-LASI (triangles). The results are very consistent with the exception of the dwell times extracted from the 5 nt sample, which was predicted with a low confidence distribution due to the low SNR and limited amount of information in the one-channel input. For comparison, the dwell times obtained from TIRF data are displayed in light gray.

Next, we applied Deep-LASI to smFRET measurements on proteins. We previously used dual-color FRET studies to probe the nucleotide-dependent conformational states^38^ of Ssc1, a mitochondrial heat-shock protein Hsp70 in yeast. By fluorescently labeling both, the nucleotide-binding domain and the substrate-binding domain, we investigated the influence of ADP on the inter-domain separation via smFRET. For the different ADP concentrations, Deep-LASI identifies the underlying FRET states in line with the manually evaluated data^38^ (Figure 6c). It correctly identifies transitions between two distinct states, a loosely docked conformation with high FRET efficiency (E = 81 %) and a separated undocked state (E = 50%), as shown in Figure 6d). The automated data analysis of Deep-LASI confirmed the ADP-dependent kinetics of the domain sensor in good agreement with previous, manually evaluated results^38^ (Figure 6e,f) and demonstrated its proficiency for unsupervised data evaluation of smFRET data on proteins.

Finally, we tested the automatic analysis of Deep-LASI applied to a different microscopy approach for smFRET, i.e. confocal single-molecule data on immobilized molecules that can be collected with microsecond time resolution. We chose the double-labeled DNA origami structure introduced in Figure 2a with various binding site combinations to tune the dwell times over different time scales. These include highly asymmetric transition probabilities as well as dwell times on the order of ~1 ms. Figure 6g shows a representative intensity trajectory of a DNA origami structure containing 6 nt complementary overhangs that was classified as ‘dynamic’ and the corresponding predictions of the state classifier. Although the unquenched state (state 2) shows a high variance in intensity, the state classifier predicts transitions with high accuracy and confidence. In the case of the 5 nt complementary overhangs, the dwell times approach 1 ms (Figure 6h) and the output confidence of the state classifier decreases significantly due to the low signal-to-noise ratio of the trace. Thus, the confidence value is an important parameter indicating the "confidence" the algorithm has in the assignment of the state and can be used as a threshold. Figure 6i (colored symbols) compares the mean dwell times extracted by Deep-LASI for all the confocal data sets with the results obtained by a newly developed shrinking-gate fluorescence correlation spectroscopy (sg-FCS) approach.^39^ For all binding site combinations with 6 nt to 7 nt complementary overhangs, dwell times obtained by both methods are in excellent agreement. The largest deviation was found for the 6 nt binding sites in the asymmetric 6 nt/7 nt sample (Figure 6i, purple) (a factor of 2) where there is large heterogeneity and limited statistics.^39^ The dwell times for the sample with 5 nt complementary overhangs follow the exponential trend observed for longer binding sites but the binning of 0.6 ms, together with the resulting low signal-to-noise ratio, reach the current limit of Deep-LASI’s state classifier. For completeness, we have included the results from Figure 4e,f in Figure 6i (gray triangles). There is a shift in dwell times between TIRF and confocal data due to the different temperatures of the two laboratories (~19 °C confocal, ~ 22 °C TIRF). Lower temperatures lead to a higher standard free energy and concomitantly longer binding time.^40,41^ In the case of the 6.5 nt binding sites sample (Figure 6i, yellow), lower dwell times are consistently observed for the TIRF data. This discrepancy is due to the difference in temporal resolution of the two measurements (2 ms for confocal vs 30 ms for TIRF). The lower temporal resolution of the TIRF measurements led to a higher probability of fast transitions being averaged out and an underestimation of the actual transition time. This is a limitation of the real experimental data and is not attributable to Deep-LASI. On the contrary, Deep-LASI can back-trace shortcomings of either technique, identify rare events, and monitor conformational changes over several time scales in an unsupervised manner.

## CONCLUSION

Deep-LASI is a deep-learning algorithm for the rapid and fully automated analysis of one-, two- and three-color single molecule assays. Employing state of the art neural network architectures optimized for time series data, we extend the classification of two-color FRET trajectories to include single- and three-color data, analyzed the photobleaching information and incorporated a full state-transition classification. Deep-LASI addresses the need for unbiased, high-throughput screening of fluorescence intensity trajectories. This opens new possibilities for single-molecule assays and enables a timely analysis of complex experimental approaches thanks to the efficient and retrainable neural network architecture of Deep-LASI. It has a high potential for applications in a myriad of fields including biotheranostics, sensing, DNA barcoding, proteomics and single-molecule protein sequencing. We envision that deep-learning approaches along with single-molecule sensitivity will dramatically assist and accelerate analytics and be indispensable in the future.

## METHODS

Methods, including statements of data availability and any associated codes and references are available in the online methods

## ONLINE MATERIAL AND METHODS

### 1. Chemicals

Chemicals were purchased from Sigma-Aldrich and used without further purification, if not stated otherwise. Chemicals include acetic acid, agarose, ammonium persulfate, (3-aminopropyl-) triethoxysilane (APTES), biotin-poly(ethylene glycol)-silane (*biotin-PEG*, MW3000, PG2-BNSL-3k, Nanocs, NY; USA), bovine serum albumin (BSA; New England Biolabs, Ipswich, Ma, USA), ‘Blue Juice’ gel loading buffer (ThermoFisher Scientific), ethylene-diamine-tetraacetic acid sodium salt dehydrate (EDTA-Na_2_ × 2H_2_O), glycerol, magnesium chloride (MgCl_2_ × 6H_2_O), 2-[methoxy(polyethyleneoxy)propyl]trimethoxy-silane (*mPEG*, #AB111226, abcr; Germany), phosphate-buffered saline (PBS), protocatechuate 3,4-dioxygenase from *Pseudomonas sp*. (PCD), protocatechuic acid (PCA), streptavidin, sodium chloride, Tris base, Tris HCl, and 6-hydroxy-2,5,7,8-tetramethylchroman-2-carboxylic acid (Trolox) and beta-mercaptoethanol (βME).

All unmodified staple strands (Supplementary Note 7, Supplementary Table SN7.2) used for DNA origami structure folding are commercially available and were purchased from Integrated DNA Technologies®. Staple strands with modifications (Supplementary Table SN7.3 and SN7.4) were obtained from Biomers (Supplementary Table SN7.3: Biotin; Supplementary Table SN7.4: Atto488) and Eurofines Genomics (Supplementary Table SN7.4: binding sites, Cy3b and Atto647N).

### 2. DNA origami structures: Assembly and Purification

Preparation of the L-shaped DNA origami structures follows the procedures described previously by Tinnefeld et al.^5,30^. In brief, the L-shaped DNA origami structures were folded with a 10-fold excess of unmodified and labeled oligonucleotides to the complimentary 8064 bp scaffold strand in folding buffer, which contained 40 mM Tris base, 20 mM acetic acid, 20 mM MgCl_2_ × 6 H_2_O, and 1 mM EDTA-Na_2_ × 2 H_2_O. For folding, a nonlinear thermal annealing ramp over 16 hours was used.^42^

After folding, the DNA origami solution was cleaned via gel electrophoresis in 50 mL 1.5% agarose-gel containing 1× gel buffer (40 mM Tris base, 20 mM acetic acid, 12 mM MgCl_2_ × 6 H_2_O, and 1 mM EDTA-Na_2_ × 2 H_2_O). The gel pockets were filled with a solution of 1× ‘Blue Juice’ gel loading buffer and the DNA origami solution. The ice-cooled gel was run for 2 h at 60 V. When samples were to be recovered from the gel, the staining step was omitted and the Cy3b fluorescence was used instead to identify the correct DNA origami structures. Gel extraction was performed via cutting with a scalpel and squeezing the gel with a Parafilm® (Bernis®) wrapped glass slide. The concentration was determined by absorption spectroscopy on a NanoDrop 2000 device (ThermoFisher Scientific). Purified DNA origami structures were kept in storage buffer, i.e., in 1x TAE buffer (40 mM Tris base, 20 mM acetic acid and 1 mM EDTA-Na_2_ × 2H_2_O) with 12.5 mM MgCl_2_ × 6 H_2_O.

### 3. Sample preparation for multicolor prism-type TIRF experiments

Labeled DNA origami molecules were immobilized in flow channels formed between a coverslip and a surface functionalized quartz prism. The surfaces were sandwiched on top of each other and sealed by a molten, pre-cut Nesco film (Nesco) channel. The employed prism surface was functionalized before with a biotin-PEG/mPEG coating to achieve surface passivation and prevent unspecific binding. Before the TIRF experiments, the prisms were first flushed with PBS and then incubated with a streptavidin solution (0.2 mg/mL) for 15 min. Afterwards, the sample holder was washed 3× with PBS to remove free streptavidin and then with storage buffer (1x TAE, 12.5 mM MgCl2, pH=8.4). Next, the DNA origami sample was diluted to 40 pM in storage buffer, added to the flow chamber and immobilized to the prism surface via the biotin-streptavidin linkage. After 5 min, untethered DNA origami structures were removed by rinsing the chamber 3× with storage buffer. Next, the attached fluorophores on the DNA origami structure were photostabilized by a combination of ROXS and an oxygen scavenging system^43^. For this, the sample chamber was flushed with photostabilizing buffer and finally sealed for enzymatic oxygen depletion before starting the prism-type TIRF experiment. The photostabilization buffer was mixed from two solutions in a ratio of 50:1: the (1) storage buffer containing 1 mM aged Trolox and 1 mM PCA, and (2) a 50x stock solution containing PCD (10 μM PCD, 50% glycerol, 50 mM KCl, 143 mM βME, 100 mM Tris HCl, 1 mM EDTA-Na_2_ × 2H_2_O).

All two- and three-color FRET experiments were carried out using msALEX^31^, i.e., two- or three excitation lasers were alternated frame-wise. The lasers of different excitation wavelengths were synchronized using an acousto-optical filter (OPTO-ELECTRONIC, France) with the camera frame rate using an FPGA that synchronizes the excitation and simultaneous detection on the EMCCD cameras at 50 ms exposure for 2000 (two-color) and 2400 (three-color) frames. The laser powers were set to 28 mW (0.022 mm^2^, 491 nm), 16 mW (0.040 mm^2^, 561 nm) and 10 mW (0.022 mm^2^, 640 nm) for B-Y-R excitation.

### 4. Multi-color TIRF setup

Single-pair FRET experiments on surface immobilized DNA origami structures were carried out on a home-built TIRF microscope with prism-type excitation as previously published^44^. Three laser sources (Cobolt, Solna; Sweden) at 491 nm, 561 nm, and 640 nm are available, and used for triple-color TIRF experiments with an alternation rate of 27 Hz (including a 2.2 ms frame transfer rate) between the B-Y-R laser excitation. The resulting emission was collected by a 60× water immersion objective (60×/1.27 WI Plan Apo IR, Nikon), cleaned up with a notch filter (Stopline^®^ Notch 488/647, AHF) and the red emission was separated from the blue/yellow emission by a dichroic mirror (630DCXR AHF; Germany) followed by separation of the blue and yellow emission (560DCXR AHF). The emission was spectrally filtered (AHF Analysentechnik, Tübingen, Germany) for the blue (ET525/50), yellow (HQ595/50) and red (ET685/40) collection channels and afterwards detected on three EMCCD cameras (Andor iXon (1×)/iXon Ultra (2×), Andor Technologies, Belfast; UK) via the supplier’s software Andor Solis. Synchronization and alternation of the exciting laser sources as well as the frame-wise data acquisition on three separate cameras was achieved using a LabView-written program that controls a field programmable gate array (FPGA). While the program starts the measurement, the FPGA synchronizes the execution of the hardware via TTL pulses, i.e., it controls switching on/off the excitation sources by direct modulation of the AOTF (491, 561, 640 nm), while simultaneously starting the data acquisition by the three cameras. The videos were analyzed afterward by a custom-written MATLAB program (*Mathworks*, Massachusetts, USA).

### 5. Single-molecule data analysis

Time traces of individual, fluorescently labeled DNA origami structures were extracted from measurements using one, two or three cameras for one, two and three-color experiments respectively using a custom-written MATLAB program (*Mathworks*, Massachusetts, USA). All raw data were recorded by EMCCD cameras (i.e., frames with 512 × 512 pixels containing fluorescence intensity information) and stored as TIFF stacks. The resulting traces are then either analyzed using Deep-LASI (Supplementary Notes 2-3) or manually (Supplementary Note 5). The regions of single molecule traces that represent single molecules with photoactive fluorophores were selected and the resulting states and kinetics were determined.

## DATA AVAILABILITY

Data sets will be available from Zenodo [DOI].

## CODE AVAILABILITY

A version of the program is available on GitHub [link]. The source code implemented are available from the corresponding authors upon reasonable request.

## ACKNOWLEDGEMENT

We thank Julian Heeg for helping with data collection. We thankfully acknowledge the financial support of the Deutsche Forschungsgemeinschaft (DFG, German Research Foundation) – Project-ID 201269156 – SFB 1032 Project B03 (to D.C.L.), Project A13 (to P.T.), individual grants to PL696/4-1 (to E.P.), TI 329/9-2, project number 267681426, TI 329/14-1, TI 329/15-1 (to P.T.) and Germany’s Excellence Strategy − EXC 089/1−390776260. D.C.L. and P.T. gratefully acknowledge funding from the Federal Ministry of Education and Research (BMBF) and the Free State of Bavaria under the Excellence Strategy of the Federal Government and the Länder through the ONE MUNICH Project Munich Multiscale Biofabrication. P.T. acknowledges the support of BMBF (SIBOF, 03VP03891). D.C.L., P.T. and E.P gratefully acknowledge the financial support of the Ludwig-Maximilians-Universität München via the Department of Chemistry, the Center for NanoScience (CeNS) and the LMUinnovativ initiative BioImaging Network (BIN).

## AUTHOR CONTRIBUTIONS

S.W. developed and implemented the deep-learning algorithm Deep-LASI and performed the Deep-learning assisted analyses, J.B. prepared the DNA origami samples under the supervision of P.T., P.A. collected the single-molecule TIRF data, P.A. and E.P. performed the manual analysis of the smFRET data, C.B.S. wrote the manual analysis software in which Deep-LASI was incorporated, S.W. and E.P. wrote the first draft of the manuscript, S.W. and E.P. designed the figures, all authors contributed to revising the manuscript. E.P. and D.C.L. supervised the project.

## Supplementary Information

### SUPPLEMENTARY NOTE 1: NEURAL NETWORK

#### 1.1. Architecture

For the Deep-LASI software package, two different neural-network architectures are used. One architecture is for trace classification and another for the number of states and state transition classification (Supplementary Figure SN1.1). Both architectures are hybrids of a convolutional neural network (CNN) and a long short-term memory (LSTM) model, which were designed using TensorFlow with Keras API.^1^ The CNN framework was inspired by an Omni-scale 1D-CNN, which elegantly solves the problem of finding the optimal kernel sizes by making it part of the training process.^2^ For all of our presented tasks, the omni-scale CNN LSTM hybrid architecture outperformed pure LSTM or ResNet^3^ models. We did not employ the full range of prime numbers suggested for the kernel sizes as we found the accuracy did not increase above 23. Hence, the number of trainable parameters was greatly reduced. In the trace classifier model, we added a 1×1 convolution layer for dimensionality reduction to further increase efficiency without a trade-off in validation accuracy. We omitted any kind of pooling and averaging layers as they significantly decrease the accuracy. All convolution layers are trained using the follow conditions: bias, same padding, a stride of 1, and a He Normal distribution^4^ for the kernel initializer. Each convolution layer is followed by a batch normalization layer and an activation layer using the rectified linear unit (ReLU) activation function.^5^

**Figure SN1.1:**
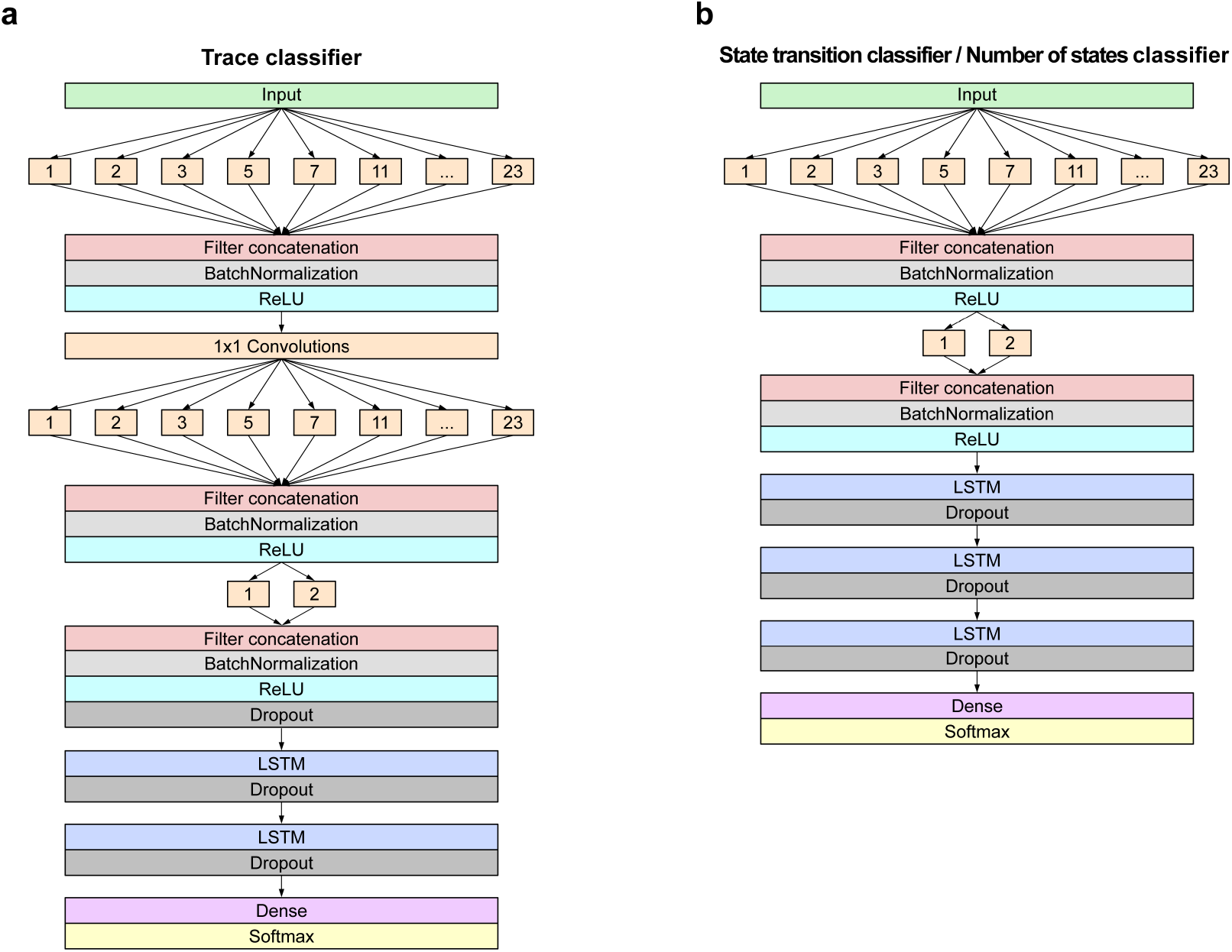
The deep neural network architectures used for the trace classifier **(a)**, and for the state transition classifier and the number of states classifier **(b)**.

#### 1.2. Trace classifier architecture

The trace classifier consists of four convolution layers followed by two LSTM layers and one fully connected layer as the feature extraction module (Supplementary Figure SN1.1a). In the first convolution layer, the input is fed into 10 layers with 32 filters each. The kernel filter size is varied between layers with sizes given by the prime numbers from 1 to 23. All layers are stacked sequentially, i.e., they operate on the same level of depth. The second convolution layer serves as a dimensionality reduction layer with 32 filters and a kernel size of 1. The third convolution layer has the same hyperparameters as the first layer. A fourth convolution layer is added, composed of two branches with 32 filters each and kernel sizes of one and two, which allows the receptive fields of the network to cover all possible integers. The output of the CNN is fed into a LSTM layer with 128 units, followed by a second LSTM layer with 32 units and the final dense layer for classification. For the training procedure, we placed a dropout layer at a rate of 0.22 before the first LSTM layer and two dropout layers at a rate of 0.5 after the two consecutive LSTM layers to maximize the validation accuracy and reduce overfitting.

#### 1.3. State transition classifier and number of states classifier architecture

The main difference in architecture between the transition classifier and the trace classifier are the depths and widths of the CNN and LSTM structure. The state transition classifier is composed of two convolution layers, three LSTM layers, and one final dense layer (Supplementary Figure SN1.1b). The kernel sizes of the first convolution layer are prime numbers in the range of 1 to 23 with 64 filters each. The second convolution layer has kernel sizes of one and two with 32 filters each. The CNN substructure is directly followed by three LSTM layers with 128 units. Dropout layers are placed after each LSTM layer using a rate of 0.5. At the end, a fully connected layer (or dense layer) is used to reduce the output of the network into the number of given categories.

### SUPPLEMENTARY NOTE 2: TRAINING

#### 2.1. Training procedure

Both the trace classifier and transition classifier models were trained using the Adam optimizer with the default settings^6^. We used a hybrid method of increasing the batch size and lowering the learning rate during training. The entire training set of ca 200,000 traces is feed into the neural network in batch sizes of 32 traces until the network has seen all traces (referred to as an epoch). After the network has seen all traces once, some input units are randomly set to 0 using dropout layers and the dataset is fed again in batches to the neural network in the next epoch. The dropout layers reduce overfitting and allow generalization of the learned information. When the validation loss is not significantly lowered within 4 epochs, the batch size is doubled. An initial learning rate of 0.001 was decreased analogously by factors of 10 after a batch size of 512 was reached.

#### 2.2. Training data set preparation

To generate training datasets, we found the approach of using simulations, originally described in Thomsen et al.^7^, to be the most promising. This is especially true for three-color models capable of detecting state transitions or photobleaching events of each dye individually. A manual collection of labeled traces on a scale large enough for adequate training would be prone to biases and/or errors due to incorrect trace identification. In addition, the datasets would not be optimized for microscope setups with different characteristics. The signal and noise characteristics of smFRET data is well enough understood that simulated data can accurately reproduce the characteristics of real data. Beside the architecture itself, the main differences between our trace classifiers and the DeepFRET model^7^ is the ability to classify one-color and three-color data and to predict the photobleached frames of each fluorophore separately. We adopted the categories ‘dynamic’, ‘static’, ‘noisy’, and ‘aggregate’ while implementing additional categories for all possible photobleaching events. The ‘artifact’ category includes false localizations, overestimated background and random perturbations of the intensity traces.

For the simulated data, idealized intensity traces for each primary category (i.e., non-photobleaching category, ‘dynamic’, ‘static’, ‘noisy’, ‘aggregate’ and ‘artifact’) are generated and then photobleaching steps are added for the different dyes by randomly determining the survival time of the dye from a given exponential distribution. The addition of photobleaching as well as other processes in the simulation may lead to alterations in the label of the given trace. For example, a simulated ‘dynamic’ trace that does not undergo a transition before photobleaching or by the end of the trace would be recategorized as ‘static’. To ensure that the network sees the same number of frames for the different categories including all the photobleached categories, the number of traces selected for the training set needs to be balanced. Hence, we begin by simulating ~250,000 traces of 500 frames for each primary category. After including photobleaching, the number of labeled frames for each category is determined. The category with the minimum number of frames determines a threshold at which additional traces, depending on their present classification, are added or excluded from the final training dataset.

The typical number of traces included in the final training dataset is approximately 200,000. This balancing procedure ensures that no category is over- or underrepresented across all frames and minimizes biases of the trained deep neural networks. Supplementary Figure SN2.1 shows the cumulative distribution of category labels in each training set used for trace classification.

**Figure SN2.1:**
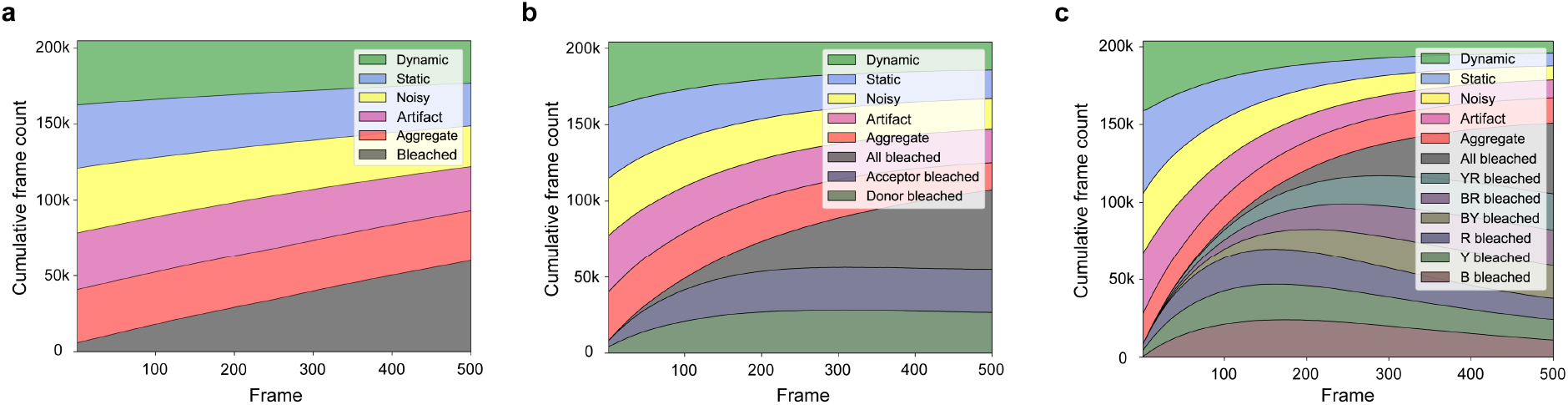
Cumulative distribution of labeled categories in training data sets for one-color **(a)**, two-color **(b)** and three-color data **(c)**. The same dataset was used for training the continuous wave two-color network as for the two-color ALEX network with the exception that the ALEX channel was not included.

For training of the state transition network, only frames where all dyes are photoactive are included and hence photobleaching can be ignored for training this network. The number of categories in the training sets then equals the number of states in the model. The visible states are first counted and sorted according to their chronological order before the state of each frame is assigned. This results in the first observed state always receiving the first label regardless of the FRET efficiency value and hence a state label only corresponds to a particular FRET value when a given dataset is analyzed globally.

#### 2.3. Simulation of single molecule traces

Single-molecule intensities traces are simulated by first initializing the number of traces to be simulated (250,000 in our case) and the probability of the trace being a single-molecule trajectory or an ‘aggregate’. All parameters used for simulating single-molecule traces are given in Supplementary Table SN2.1. An idealized FRET efficiency trajectory is then generated in one, two or three colors where the FRET efficiency (or efficiencies) is randomly selected between 0.01 and 0.99. For single molecule trajectories, the number of states in the trajectory is selected with a probability of containing only a single state (45 % in our case for ‘static’ traces) and the remaining probability (55 % here) is equally distributed between two, three and four states (‘dynamic’). In the case of ‘dynamic’ traces, the FRET efficiency or efficiencies of each state are randomly selected from a uniform distribution between 0.01 and 0.99. The difference in FRET efficiency for the different states has to be above a given threshold (0.1 in our cases). If this is not the case, new FRET efficiencies are randomly selected until this criterion is fulfilled. The transition rates are generated by taking the inverse of the dwell time to exposure ratio drawn from a uniform distribution between 1 and 100. The generated transition matrices are then, in general, non-symmetric. Hence, we use the transition rate matrix to calculate the probability of which state is observed first. We do this by using the least-squares solution to the matrix equation *Ax* = *b*, where *A* is the transition matrix and *x* is the probabilities for observing the different states. While the calculation of the state equilibrium is not mandatory for the classification accuracy, it ensures that the output matches the ground truth input of a defined transition matrix, which was used for benchmarking the transition classifier. Once the initial state has been selected, the parameters are fed into an HMM routine^8^ and a state trajectory is generated. ‘Aggregate’ traces are always assumed to be static (uncorrelated dynamics are categorized as ‘artifact’), but the number of dye-pairs is generated from a Poisson distribution with randomly selected FRET efficiencies between each pair. Next, the idealized FRET efficiency trace is converted into normalized fluorescence intensity traces for the donor and acceptor molecules based on the FRET efficiency (discussed in more detail below). Next, photobleaching of the fluorophores are included into the trajectories. The frame at which each fluorophore photobleaches is randomly drawn from an exponential distribution. Upon photobleaching, the affected channel intensities are either set to 0 for both channels for donor photobleaching, 1 for the donor intensity upon acceptor photobleaching or recalculated using the two-color FRET equations (for three-color simulations). Blinking is then added to a fraction of the traces where each dye has a probability of being in a short-lived dark state (Supplementary Table SN2.1). At this point, ‘artifact’ traces are generated from ‘static’ or ‘dynamic’ traces with a given probability by subtracting a constant from the trace (to simulate overestimation of the background correction), adding random fluctuations to the total intensity (to simulate among other things new molecules or aggregates flowing through the observation volume or simulating molecules in the background mask), flipping the traces (to simulate molecules that turn on during the experiment) and/or adding non-correlated signal in the different channels. To account for non-uniform brightness of the individual molecules, all excitation channels are multiplied by a scaling factor that is randomly selected from uniform distributions. In particular, the red channel after red excitation, *I*_RR_, can reach scaling factors up to three times higher than the other two channels. This allows the trace classifier to correctly analyze data sets in which high red laser powers were used to increase statistics for the calculation of correction factors and for making sure only a single red fluorophore is present. Without intensity scaling, the trace classifier strongly favors the aggregate category for traces with imperfect stoichiometry even when no second bleach step is present. With a given probability, additional small fluctuations in the total intensity are also added to the traces to simulate experimentally observed system instabilities (assuming sinusoidal oscillations of randomly determined frequency and amplitude). Next, we incorporate spectral crosstalk, direct excitation and differences in detection efficiency into the data by randomly selecting the respective parameters from a uniform distribution (see Supplementary Table SN2.1). In the last step, we add two or three types of random noise to the traces. The first component is intensity-independent background noise drawn from a Poisson distribution. The second component considers intensity-dependent noise contributions (i.e. shot-noise) by drawing values from Gaussian distributions. There are different descriptions of how to treat noise from EMCCD cameras. According to Basden et al., the variance in shot noise due to the EM gain is increased by a factor of two.^9^ This corresponds to a rescaling of the Gaussian distribution mentioned above. Hirsch et al describe the additional noise from EM-CCD cameras using a gamma distribution.^10^ Hence, with a given probability, we also add a third component to the noise modeled using a gamma distribution with random amplitude. After adding noise to the trajectories, we then recategorize traces with high noise as ‘noisy’. This is done by recalculating the FRET efficiency trace or traces from the intensity data. When the standard deviation for static traces or individual states of a dynamic trace are above the given threshold (Supplementary Table SN2.1, we used 0.25), the trace is categorized as ‘noisy’. Finally, the classification of the individual traces is checked and, if necessary (for example a dynamic trace that photobleaches before a transition is observed), recategorized. The dataset is then balanced, as discussed above, each trace normalized to its maximum value and then used for training.

For one-color traces, we simulated the intensity of the donor molecule although, for a single channel, it does not make a difference. The donor intensity (we refer to it as *YY* here) is given by:

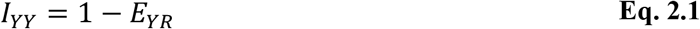

Since only one dye and one channel is observed, there is only one photobleaching category and no correction factors are included. The photoactive state of the acceptor molecule is still calculated and its influence on the donor intensity incorporated into the trace. For calculation of ‘aggregates’, fluorescent dye-pairs are added to the trace but only the donor signal is considered. Furthermore, the amount of noise is not quantified by the standard deviation of the FRET efficiency but by the signal-to-noise ratio of the channel intensity, which is defined as:

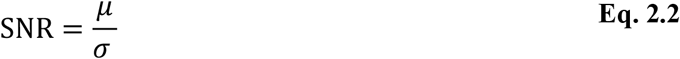

where *μ* is the mean signal intensity of the observed state and *σ* is its standard deviation. When the signal-to-noise ratio falls below the given threshold, the trace is classified as ‘noisy’.

For two-color FRET simulations, we use the same approach as described in the following section for 3-color FRET but we only consider the equations necessary for 2-color FRET, i.e., all equations including the yellow/red FRET pair.

For generating three-color FRET data, the distances and Forster radii between all three fluorophores need to be considered as they are interrelated. Assuming a minimum FRET efficiency of 0.01, the generated FRET states of the first two randomly drawn FRET pairs put constrains on the maximum possible distance for the third FRET pair. To guarantee a uniform distribution of possible FRET combinations, we randomly select two of the three FRET pairs and their corresponding FRET efficiencies. Using the two selected FRET efficiencies, a lower limited is calculated for the third FRET pair, which depends on the Förster radii of the first two FRET pairs. For example, when the yellow-red dye-pair is generated last, the dye-dye separation for *r_BY_* and *r_BR_* are calculated and then used to determine the minimum FRET efficiency (i.e., maximum separation for the third dye-pair) as given below:

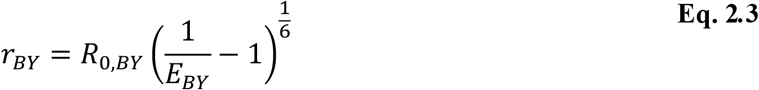

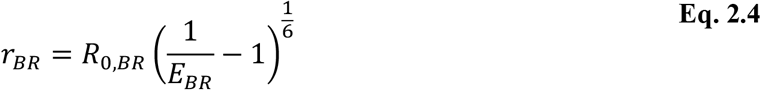

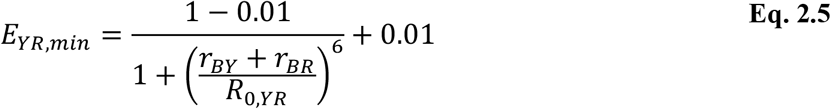

where the Forster radii *R*_0,*BY*_, *R*_0,*BR*_ and *R*_0,*YR*_ are sampled over values that are typically available using commercially available dyes pairs. The FRET efficiency *E_YR,min_* represents the lower boundary used to randomly scale the FRET trace of the yellow-red FRET pair in a correlated or anti-correlated manner. When a different dye-pair is generated last, the same equations are used where the indices are changed accordingly. For dynamic traces, this procedure is performed for all states. Once we have selected the FRET efficiencies for the different dye-pairs and states, we then convert them what would be observed for a two-color experiment. The YR dye-pair is already a two-color FRET efficiency and does not need to be corrected. When all three fluorophores are photoactive, the blue dye may be quenched by two acceptors. In this case, the distance-related FRET efficiencies *E_BY_* and *E_BR_* need to be converted into the apparent FRET efficiencies *E_BY,app_* and *E_BR,app_* via:

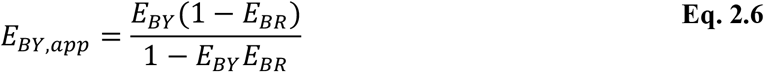

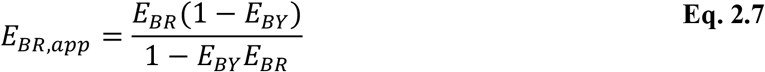

Since the input data for the neural networks are normalized, the channel intensities are initialized as follows:

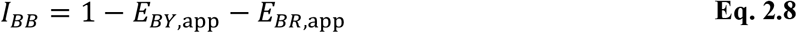

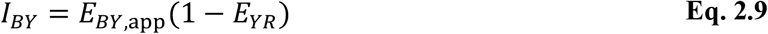

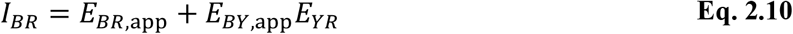

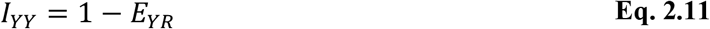

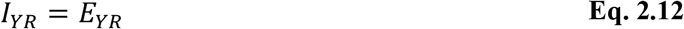

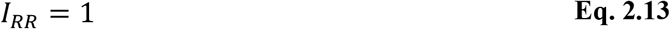

Upon photobleaching of one of the dyes in the three-color experiments, the system then reverts into the two-color case:

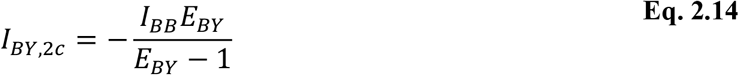

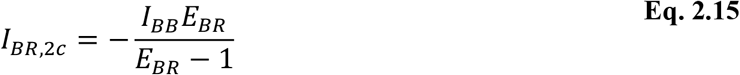

*I_BB_* is still determined by using

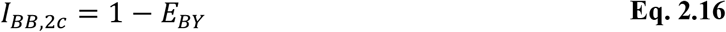

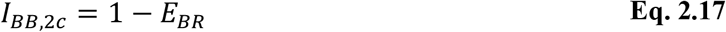

where FRET to the blinking fluorophore is set equal to zero in two-color sections of the trace. Blinking events are treated the same way as photobleaching during the frames where the one dye is off and the channel intensities are either set to 0 or recalculated using Eq. 2.16/2.17.

In three-color experiments, each channel has its own set of correction factors for differences in detection efficiency and quantum yield, *γ*, direct excitation, *de*, and spectral crosstalk, *ct*. The values are randomly drawn from a wide uniform range and implemented in the following order. First, the FRET channels are multiplied by the corresponding *γ*-factor:

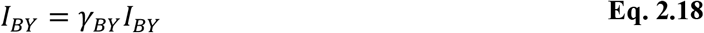

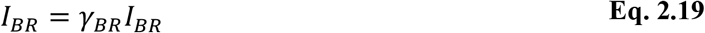

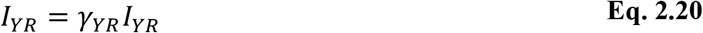

The crosstalk of the blue fluorophore leaking into the yellow and red channel after blue excitation are given by:

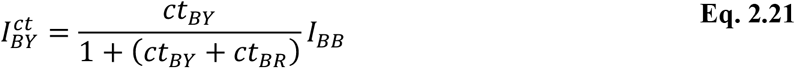

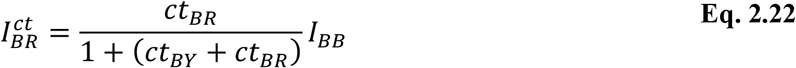

where *ct_BY_* and *ct_BR_* denote the randomly drawn crosstalk factors (Supplementary Table SN2.1). Spectral crosstalk of the yellow fluorophore into the red channel is calculated using:

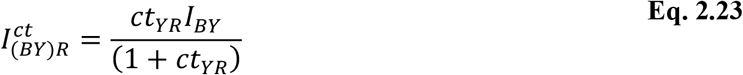

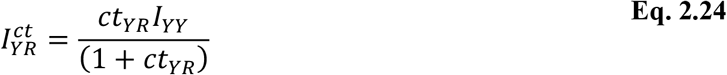

The observed intensities including all correction factors are determined by:

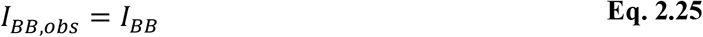

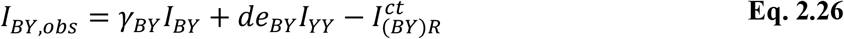

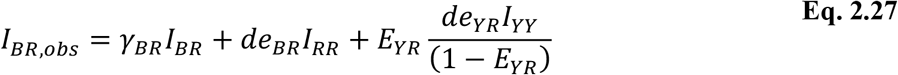

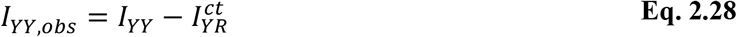

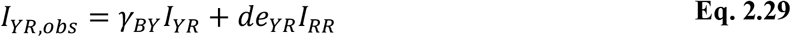

where *de_BY_*, *de_BR_* and *de_YR_* are the randomly drawn direct excitation factors from a uniform distribution (Supplementary Table SN2.1).

While the non-smFRET categories ‘noisy’ and ‘aggregate’ mimic experimental data, the category ‘artifact’ is primarily designed to increase the robustness of the trace classifier. It is important to note that the accuracy of a trained neural network to distinguish between an ‘artifact’ and any other category depends on the number of traces which are labeled as ‘artifact’ but maintain a strong resemblance to the original trace. For the goal of increasing robustness, it is therefore not desirable to achieve 100% prediction accuracy as it would be caused by too easily identifiable perturbations in the training set.

#### 2.4. Simulation settings for training the state classifier network

Sixteen pre-trained deep neural networks are provided for state classification. Four models account for the classification and segmentation of time trajectories obtained from measurements using single-channel data acquisition, two-color FRET with continuous wave excitation, two-color FRET with ALEX, and three-color FRET with ALEX. For each type of experiment (one, two and three-color), we provide three state-transition-classifiers trained on either two, three or four observed states. The state classifier networks only use traces as input that are categorized as dynamic. Hence, the training data sets only contain valid FRET traces with at least one transition, a minimum state difference of 0.1 in FRET efficiency and no photobleaching. The transition rates are generated by drawing random dwell time to exposure ratios between 1 and 100 from a uniform distribution. Traces with a state-wise FRET distribution width above 0.25 on average are excluded from the training data set. After a dynamic trace is simulated, it is labeled according to state occupancy. Here, the first observed state always receives the first label regardless of the FRET efficiency, followed by the next observed states until the maximum number of states is reached. For three-color FRET data, every transition regardless of the dye is treated as a new state. For a two-state model, transitions of one dye can be described whereas the multi-state model also considers transitions of two dyes. Thus, we have trained the network to recognize four different states. For a system with three independently moving dyes, a minimum of 9 states would be possible in one trace. Expert users can generate a corresponding training data set by setting the algorithm parameter ‘static dyes’ to ‘None’.

**Supplementary Table SN2.1.**
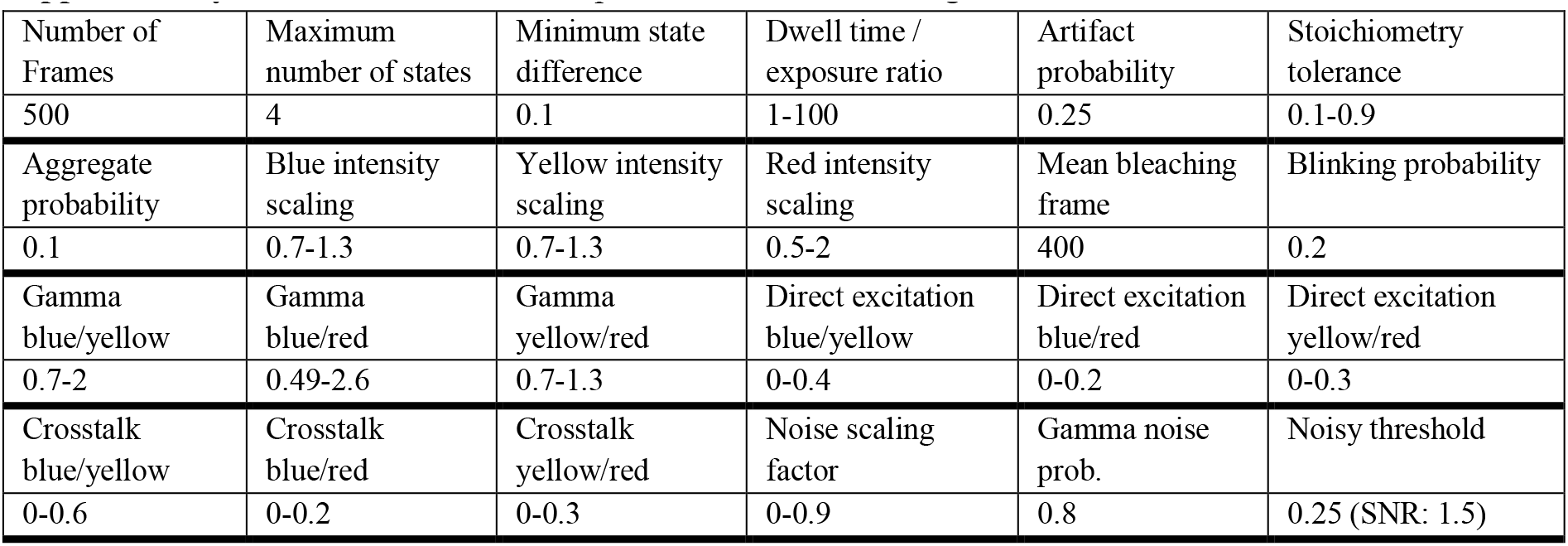
Simulation parameters for the training data sets.

| Number of Frames | Maximum number of states | Minimum state difference | Dwell time / exposure ratio | Artifact probability | Stoichiometry tolerance |
| --- | --- | --- | --- | --- | --- |
| 500 | 4 | 0.1 | 1-100 | 0.25 | 0.1-0.9 |
| Aggregate probability | Blue intensity scaling | Yellow intensity scaling | Red intensity scaling | Mean bleaching frame | Blinking probability |
| 0.1 | 0.7-1.3 | 0.7-1.3 | 0.5-2 | 400 | 0.2 |
| Gamma blue/yellow | Gamma blue/red | Gamma yellow/red | Direct excitation blue/yellow | Direct excitation blue/red | Direct excitation yellow/red |
| 0.7-2 | 0.49-2.6 | 0.7-1.3 | 0-0.4 | 0-0.2 | 0-0.3 |
| Crosstalk blue/yellow | Crosstalk blue/red | Crosstalk yellow/red | Noise scaling factor | Gamma noise prob. | Noisy threshold |
| 0-0.6 | 0-0.2 | 0-0.3 | 0-0.9 | 0.8 | 0.25 (SNR: 1.5) |

### SUPPLEMENTARY NOTE 3: TRAINING VALIDATION

In the following sections (3.1–3.3), the final validation of every deep neural network is shown via confusion matrices. Approximately 20,000 new simulated traces were generated and fed into each trained model. Each row of the confusion matrices represents the instances in a ground truth category while each column represents the instances in a predicted category. The diagonal values report the percentage of true positives and true negatives whereas the off-diagonal values are the false negatives and false positives.

#### 3.1. Trace classifiers

Confusion matrices for the trace-classifier networks are shown in Supplementary Figures SN3.1 and SN3.2. The single-channel classifier has the lowest overall performance, in particular, due to a higher rate of falsely classifying random perturbations in ‘artifact’ frames (88 % precision) and misinterpreting ‘dynamic’ traces as ‘static’ (5 % false negative rate). The two-color and three-color models achieve similar accuracies for recovering smFRET frames with at least 93 % precision in correctly predicting ‘dynamic’ frames and 96 % precision for ‘static’ frames. In general, most of the false predictions concerning smFRET categories come from the high resemblance of ‘static’ frames, ‘dynamic’ frames with low contrast between states and ‘noisy’ frames close to the defined threshold. Here, the tolerance towards noise, defined as the mean standard deviation of the observed FRET efficiencies for all states, was set to 0.25. The highest sensitivity for detecting photobleached dyes (>98 %) is achieved by ALEX-enabled models for two- and three-color data. The continuous wave models depend on the contrast in intensity between the quenched and photobleached dyes, causing a significant decrease in sensitivity down to 91 % for detecting a photobleached acceptor. However, falsely predicted ‘acceptor bleached’ frames were mostly misclassified as either ‘aggregate’ or ‘artifact’ and would still be excluded from further analysis.

In addition to the confusion matrix for all available categories, we also calculated a binary trace classifier confusion matrix where we separated the frames into those that were accepted for further analysis (i.e., from ‘static’ and ‘dynamic’ traces without photobleaching) and those that were rejected (‘photobleached’, ‘aggregate’, ‘artifact’ and ‘noisy’ traces and/or frames). All trace classifier models achieve a minimum combined precision of 97 % in predicting smFRET categories, i.e., ‘static’ or ‘dynamic’, and 96 % in predicting non-smFRET categories (Supplementary Figure SN3.1 and SN3.2).

**Figure SN3.1:**
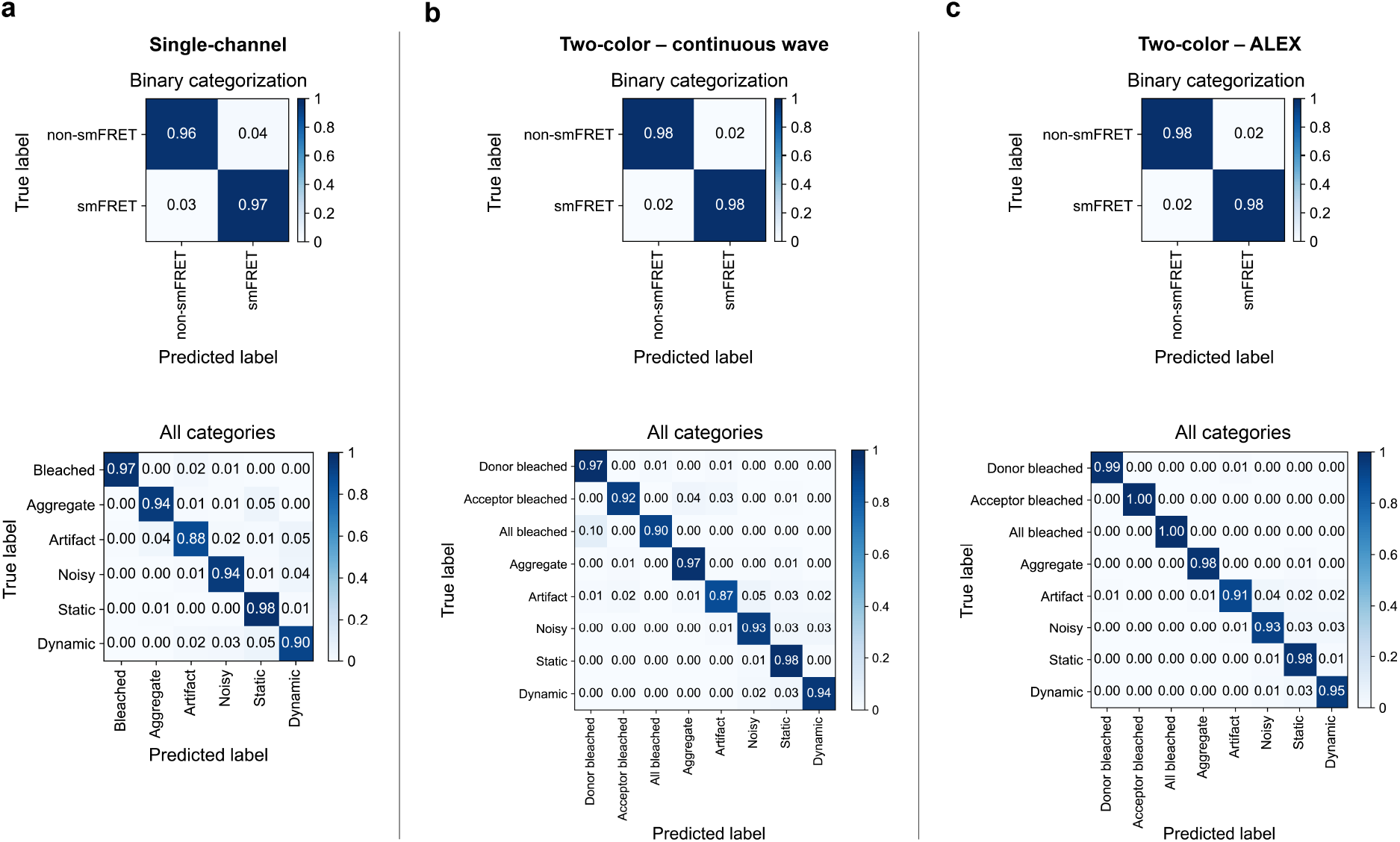
Confusion matrices for 1-color and 2-color trace classification. Prediction accuracies depicted as confusion matrices for the **(a)** single-channel, **(b)** two-color continuous wave and **(c)** two-color ALEX models. The upper panels show the binary assignments into valid smFRET and non-smFRET categories. The detailed categorization is shown in the lower panels.

**Figure SN3.2:**
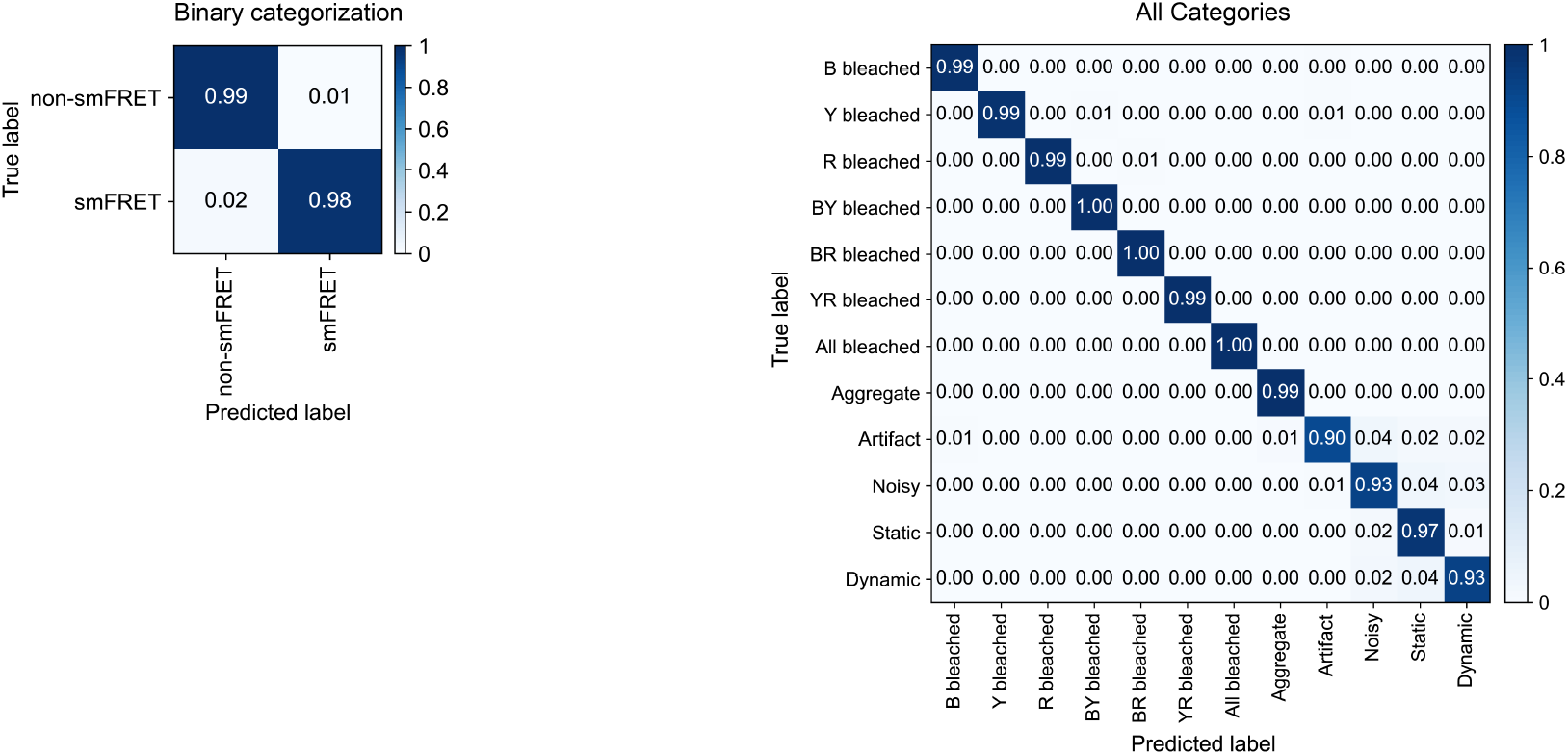
Confusion matrices for 3-color trace classification. The left panel shows the binary assignments into valid smFRET and non-smFRET categories. The detailed categorization is shown in the right panel.

#### 3.2. Number-of-states classifiers

After classifying the individual traces, the dynamics are analyzed. One option is to classify the number of states in a particular trace, i.e. to run the number of states classifier for the type of data measured. Supplementary Figure SN3.3 shows validation of the deep neural networks trained on traces containing the given number of observed states. Only traces classified as ‘dynamic’ by the trace classifiers serve as input, hence the first category is for two observed states. The category of five observed states serves as a safeguard against traces that may be out of the scope of the pretrained state transition classifiers. All models achieve a high accuracy of at least 98 % in distinguishing two-state from multi state traces. The lowest accuracies are achieved in separating four-state from five-state traces, ranging from 86 % (single-channel) to 89 % (5-channels). The overall performance increases with increased number of available channels. As only dynamic information is considered in the state classifiers, the presence of the ALEX channel, though very useful for the trace classification, is no longer relevant.

**Figure SN3.3:**
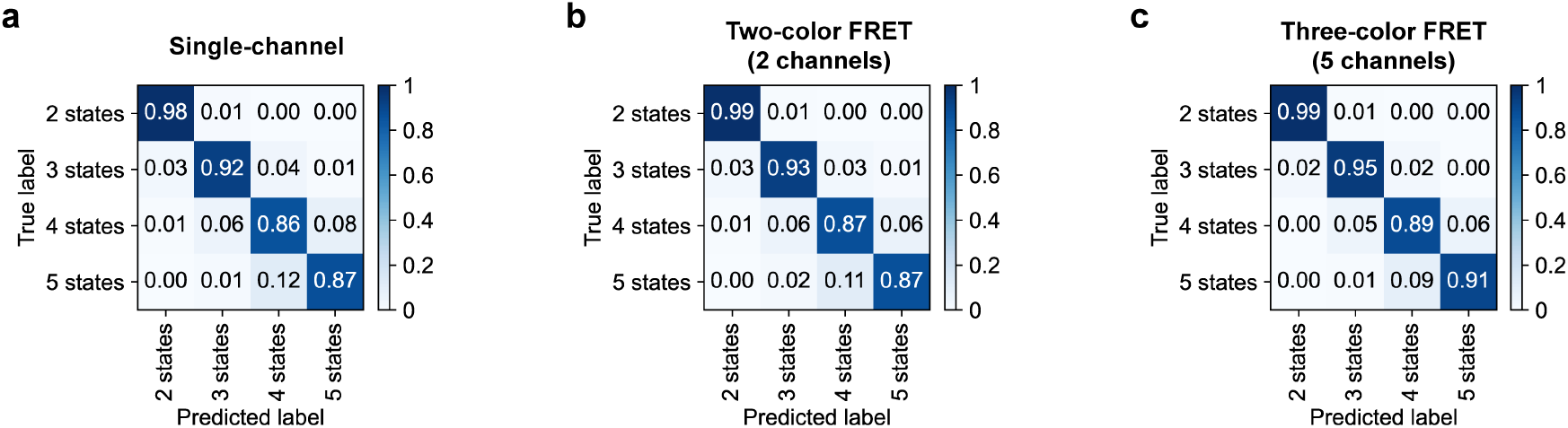
Confusion matrices for number of states classification. Confusion matrices for the **(a)** single-channel, **(b)** two-color FRET and **(c)** three-color number of states classifiers.

#### 3.3. State-transition classifiers

After estimating the number of states in a data set, the state trajectories of the individual dynamic traces are determined. This section summarizes the validation of the deep neural networks trained on the state occupancy and therefore also on the state transitions (Supplementary Figure SN3.4). The performance does not differ significantly for the two-state models with a minimum of 97 % precision for predicting the correct state and a minimum of 84 % precision for four-state systems. The performance for multi-state models increases when more channels are available

**Figure SN3.4:**
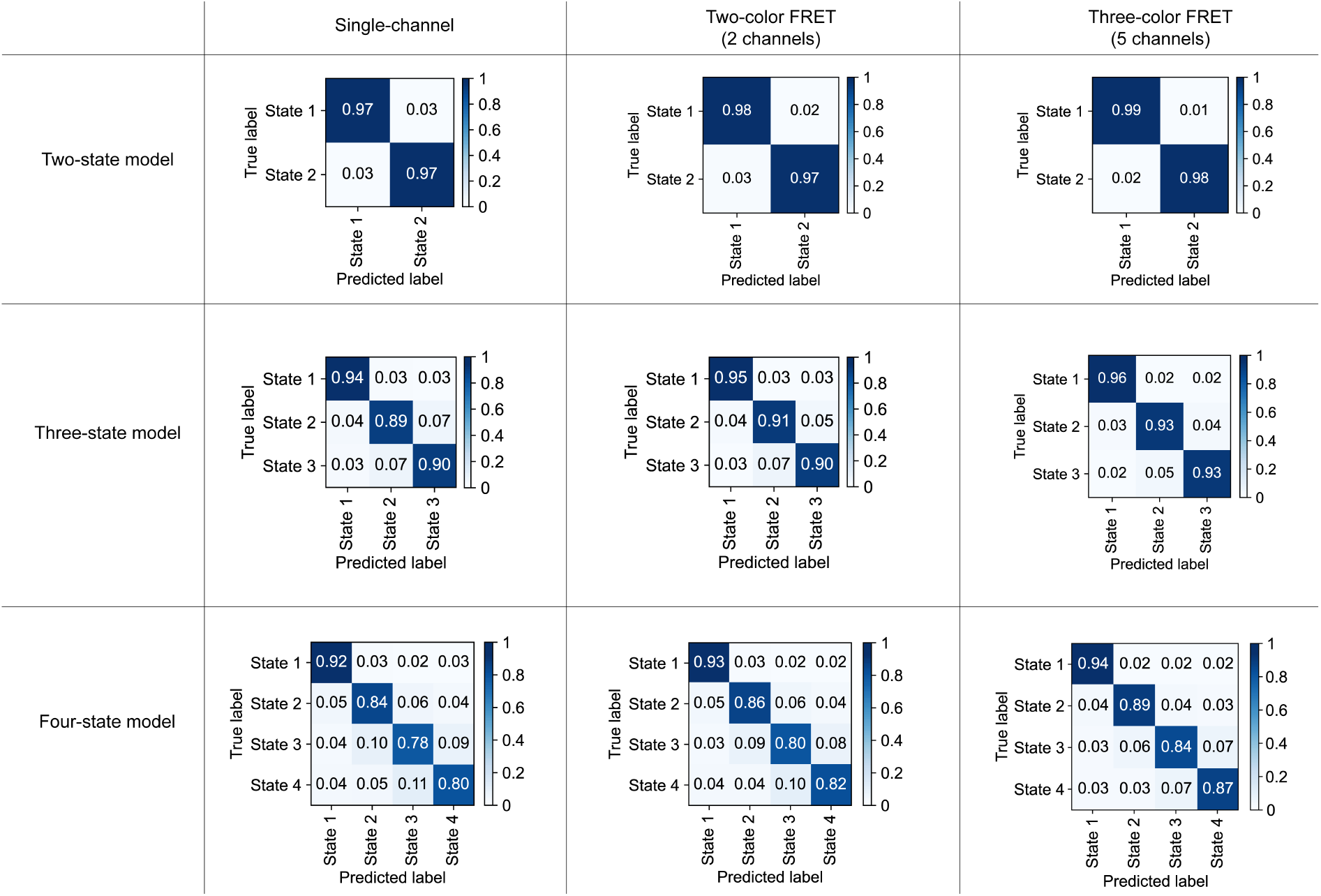
Confusion matrices for state classification. Confusion matrices for the single-channel (first column), two-color FRET (second column) and three-color state classifiers (third column) and their corresponding two-state (first row), three-state (second row) and four-state models (third row).

#### 3.4. Limitations of the state classifiers and a comparison with HMM

This section provides additional benchmarks and a comparison of the results from the state classifiers with HMM. First, we investigated how the performance of HMM and our state classifiers depends on noise (i.e. the width of the FRET distribution), difference between FRET states, the kinetic rates (dwell time to exposure ratio) and the length of observation time (number of frames) for dynamic transitions between two FRET states. For the three-color simulations, only the FRET distribution width of the yellow-red dye pair was used as the ground truth parameter to keep the continuity with two-color FRET traces and avoid averaging inconsistencies. Supplementary Figure SN3.5 shows interpolated maps of the precision of state label recovery for all models and were generated using approximately 300,000 simulated traces for each condition. The precision is the fraction of true positives divided by the sum of true positives and false positives for the state label predictions. Each map shows the precision dependency on the amount of signal noise with two of three parameters being fixed, namely the FRET state contrast (0.2), the transition rate (0.05/frame) and the number of frames (500). In general, at a fixed transition probability and number of frames (top row), the precision decreases with broader FRET distributions and smaller differences between the FRET states. All models are able to achieve a precision of at least 90% for FRET differences above 0.2 and FRET distribution widths below 0.10 with the state classifiers outperforming HMM only at high noise levels above 0.25. For a fixed contrast between FRET states (0.2) and total number of frames (500) (Supplementary Figure SN3.5, middle row), the precision of HMM remains largely independent of the dwell time to exposure ratio at a constant noise level. All DNN state classifiers show a similar overall performance but achieves a higher precision than HMM at higher noise levels for larger dwell time to exposure ratios. For fixed FRET states and kinetic rates (Supplementary Figure SN3.5, bottom row), trace length has little influence on the precision of all models below ~100 frames and the precision slightly increases for all models/classifiers above 100 frames. Again, the DNN outperform HMM at higher noise levels. In summary, while the accuracy and precision do not differ significantly between the single-channel and two-channel state classifiers, the five-channel model used for three-color FRET shows an increased performance of up to ~10 % at high noise levels. Due to the five available channels, the signal-to-noise ratio is effectively increased which leads to higher precision and accuracy as soon as the signal noise becomes a limiting factor for the other models. In addition, DNN models still predict transitions in high noise trajectories, however with decreased confidence, whereas HMM no longer finds transitions at high noise.

**Figure SN3.5:**
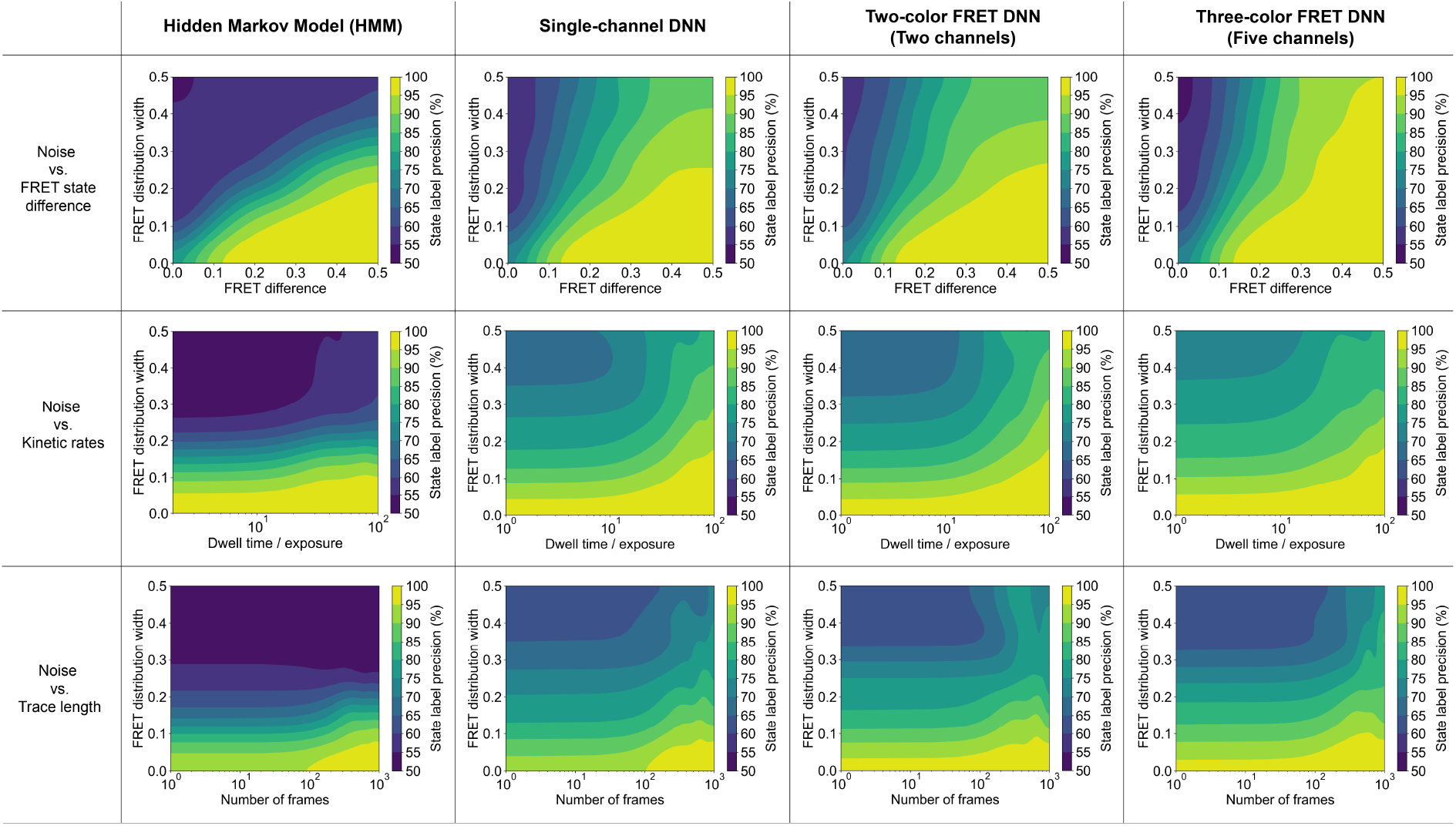
Deep-LASI state prediction compared to HMM. Precision of the state-label recovery for HMM and for the state transition classifiers as a function of noise (i.e. width of the FRET distribution), contrast between FRET states and the kinetic rates (dwell time to exposure ratio). Each map shows the precision dependency on the noise and one additional parameter: the contrast between FRET states (top row), the kinetic rate (middle row) and the number of frames (bottom row). One data set with ~300,000 traces was generated for each row with the corresponding two of the three parameters fixed (FRET efficiency for yellow/red: 0.4 and 0.6, transition probability: 0.05/frame, and number of frames: 500). The noise is defined as the mean standard deviation of the FRET signal from both states. The lower limit of the precision is set to 50 % since it represents the highest amount of uncertainty for two states.

#### 3.5. Training and validation loss

The training and validation loss for all pre-trained deep neural networks are shown in Supplementary Figure SN3.6. Of the ~200,000 traces generated for training, ~160,000 were used in each epoch for training and then the capacity of the network to generalize what it learned was tested with the remaining ~40,000 traces. The error was calculated using the categorial entropy, i.e. the loss function:

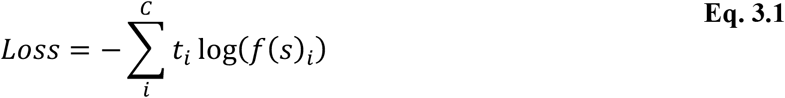

where *C* is the total number of classes, *t* is the target vector and *f*(*s*) is the one-hot encoded vector of scores. During training of the network, the loss should decrease but maintain similar values for the training data set as for the validation set. If a lower loss is observed for the training data set then for the validation data set, the network is overfitting (i.e. it is memorizing the traces rather then learning the features of the categories). All models show no or a minimal amount of overfitting. While the full number of epochs are displayed in each plot, the model with the lowest validation loss and lowest amount of overfitting was saved and implemented (indicated with an arrow).

**Figure SN3.6:**
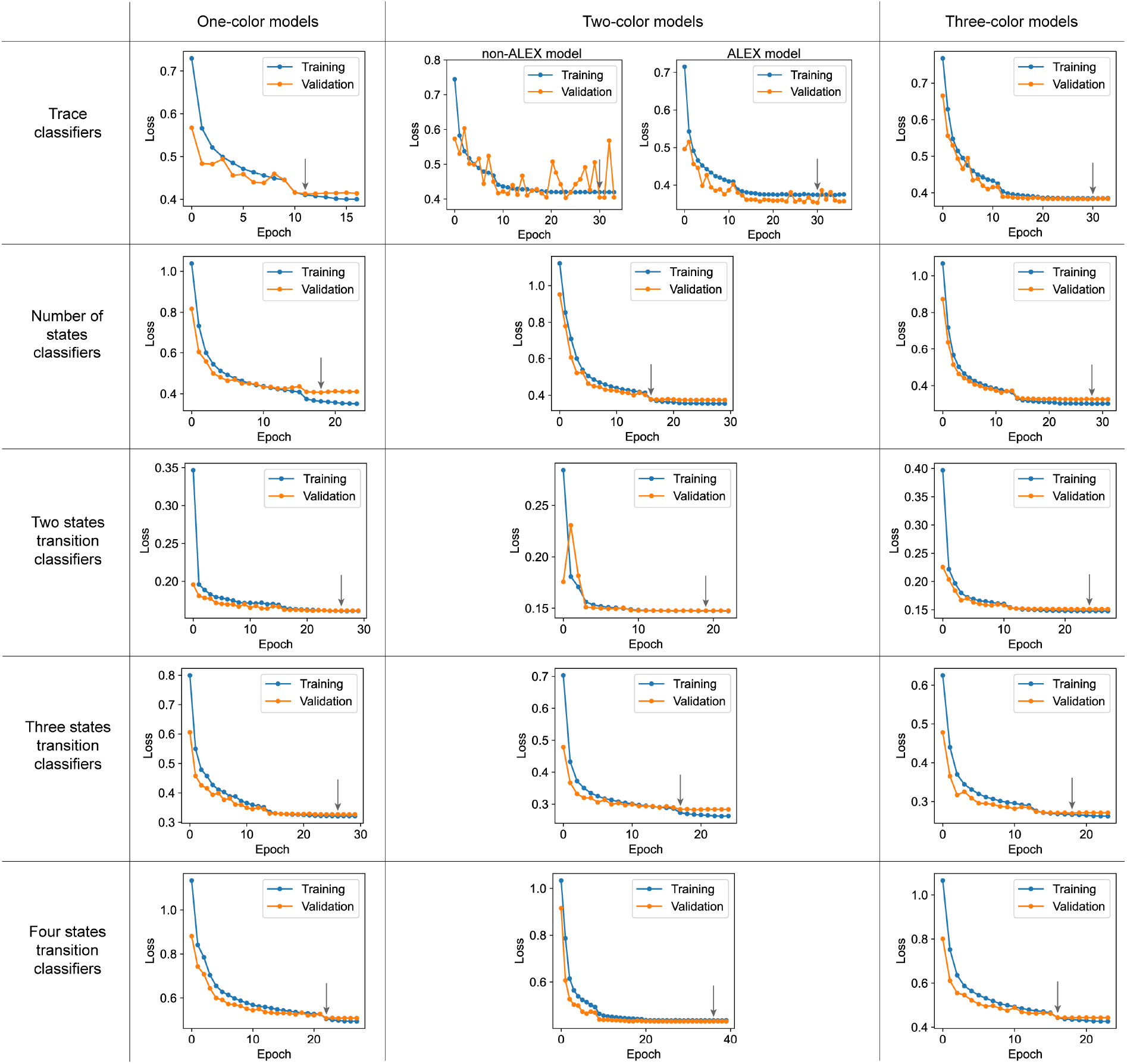
Training and validation loss of all Deep-LASI models. Each row refers to the type of classifier and each column refers to the corresponding data type. For two-color data, there are two trace classifier models, one for ALEX and a second for non-ALEX measurements. Black arrows mark the saved model used when following epochs did not decrease the validation loss and indicated overfitting.

### SUPPLEMENTARY NOTE 4: DEEP-LASI VERSUS MANUAL ANALYSES FOR THREE-COLOR DATA

We emphasize the importance of also using experimental data for testing deep learning methods trained on synthetic data since the simulations used for validation are usually generated by the same algorithm as the training data set. Deep neural networks can easily learn biases of any kind in the training data, which may have no relationship to the respective category under new conditions. Hence, the prediction of categories with respect to ground truth simulations can produce high accuracies, which may not be directly translatable to real-world examples. Therefore, we compared the performance of our network models on real data with that of experts who manually analyzed the same dataset.

We benchmarked the three-color performance of Deep-LASI by comparing the automated analysis with traces manually selected by an expert user (Supplementary Figure SN4.1). We used the three-color L-shaped DNA origami structure with two binding locations spaced at 6 and 12 o’clock with complementary binding regions of 7.5 nt (Figure 4). Deep-LASI yielded 581 usable smFRET traces versus 694 for manual selection out of a total of 2545 extracted traces (Supplementary Figure SN4.1a). The two uncorrected, framewise smFRET histograms are almost identical. The automatically extracted FRET correction factors, which is based on the predictions of the three-color trace classifier, were compared to those determined manually. The expert uses selected the relevant regions of the traces for determining various FRET correction factors by hand. Very similar distributions and median values were obtained for the YR correction factors (Supplementary Figure SN4.1b). For BR, both direction excitation and spectral crosstalk terms are small. Hence, small differences are not significant here. Due to the high stability of the yellow fluorophore, it is challenging to collect enough statistics to directly derive the detection correction factor. Hence, it is calculated from the product of the BY and YR γ factors. For BY, the distributions for spectral crosstalk from Deep-LASI and manual selection are consistent. However, for both direct excitation and the detection correction factor, there are differences of ~ 15 %. Manual selection with the blue fluorophore is difficult because of the low fluorescence intensity of the blue dye. In the manually selected regions for direct excitation, a second population is visible due to difficulties of distinguishing between a Y only fluorophore and a dim B fluorophore undergoing high FRET. Similarly, there are differences in the detection correction factor distribution. As Deep-LASI has more flexibility in choosing relevant regions of the traces for determining the correction factors, it is most likely that Deep-LASI is more accurate in these cases. FRET correction-factor determination is a potential source of human bias in the analysis of smFRET data and we demonstrate here an advantage of using a well-trained neural network for automated analysis.

**Figure SN4.1:**
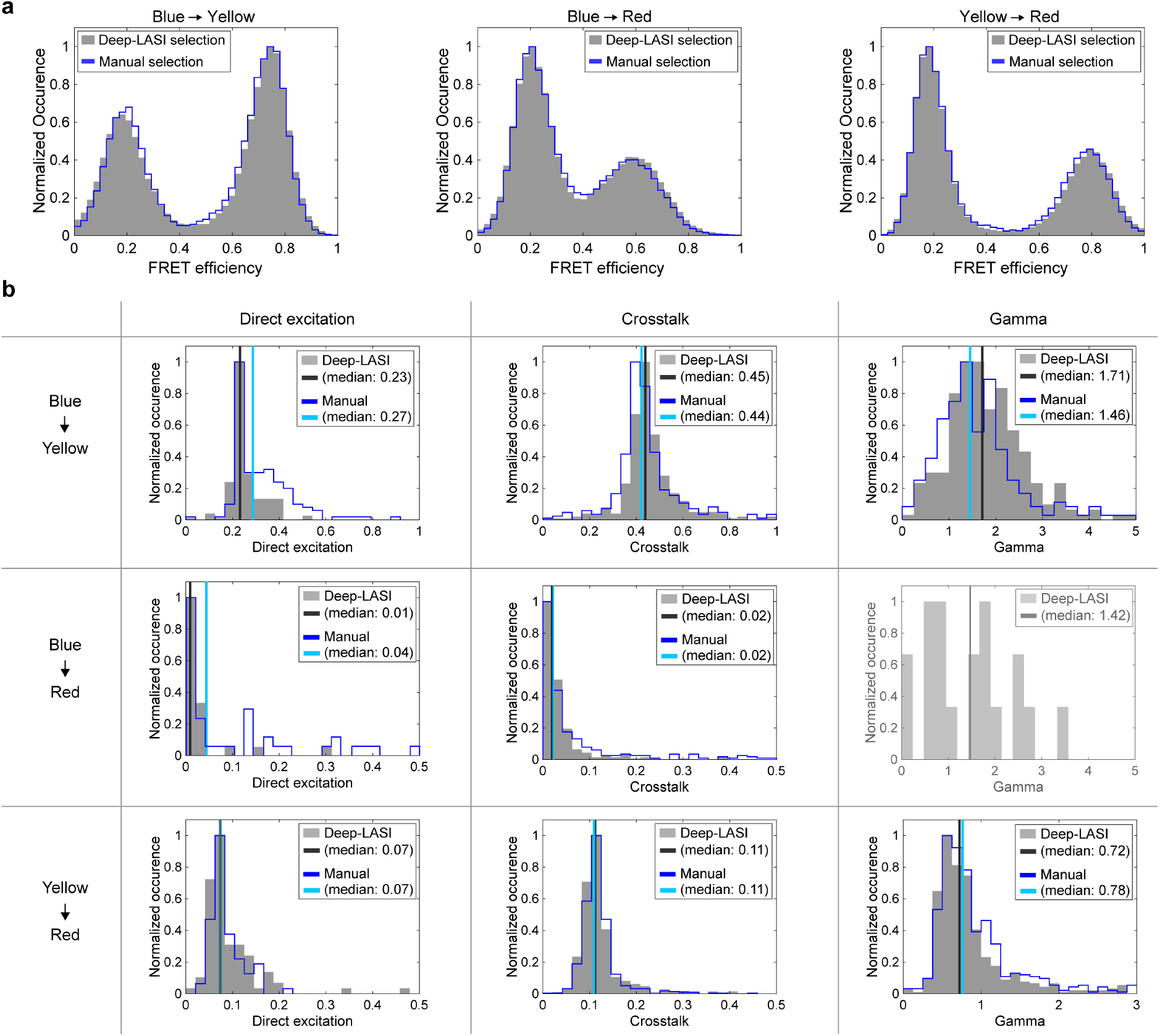
Uncorrected smFRET histograms and correction factors extracted by Deep-LASI for the 3-color 2-state DNA origami. **(a)** Uncorrected framewise smFRET histograms for BY, BR and YR calculated from traces selected manually (n=694, blue line) versus the histograms determined by Deep-LASI (n=581, gray histograms). There is excellent correspondence between the histograms. **(b)** Each panel displays the normalized distribution of available correction factors from all traces categorized as ‘dynamic’ by Deep-LASI (gray filled histograms) or manually labeled as dynamic (blue histogram line). Due to the high stability of the yellow dye compared to the blue and red dyes, the number of usable traces to calculate blue/red detection correction factor was below 5 and, hence, not determinable. Therefore, we used the theoretical value of 1.23 (for Deep-LASI compared to 1.15 for manual selection) for the blue/red gamma factor determined from the product of the gamma factors for blue/yellow and yellow/red.

### SUPPLEMENTARY NOTE 5: MANUAL ANALYSIS OF SINGLE-MOLECULE TIRF DATA

#### 5.1. Work-flow

We benchmarked the performance of Deep-LASI by comparing it to manually analyzed single-molecule data from an expert user. Starting from individual movies, the procedure for extracting the intensity information over time is highlighted in Supplementary Figure SN5.1.

The procedure begins with:

(1) a pixel-wise mapping of the position between two or three cameras for two- and three-color experiments,
(2) camera-wise localization and excitation-cycle dependent assignment of intensities, and
(3) extraction of intensities and background correction for each detection channel.

**Figure SN5.1:**
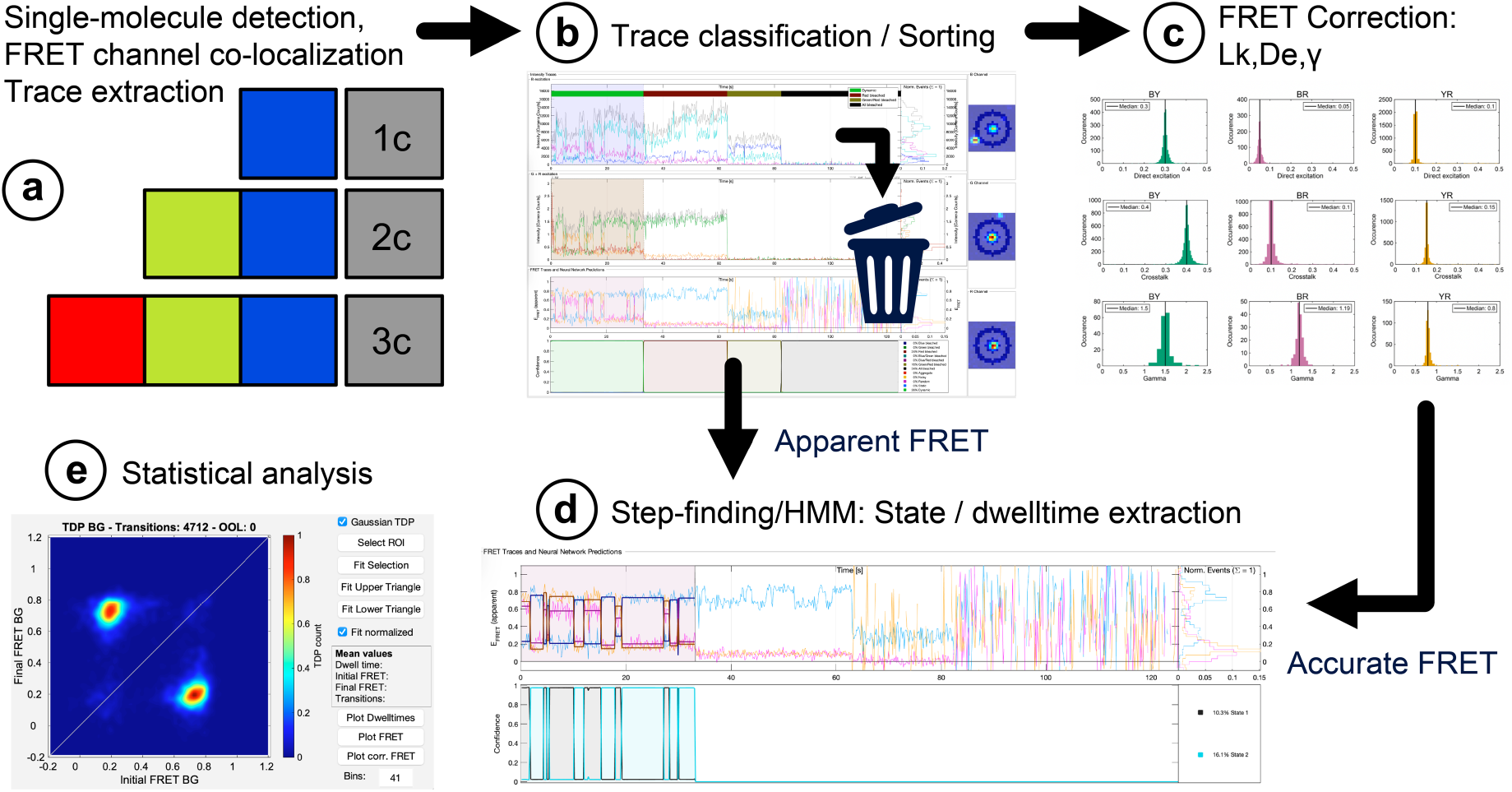
Work-flow of data extraction, sorting, analysis and evaluation. a) One-, two-, or three-color data is collected with various excitation schemes and the time-dependent intensity traces extracted and corrected for background. b) The traces are then visually inspected and sorted either for further analysis or marked as junk. Regions of the trace can be selected for smFRET evaluation or for correction factor determination. c) After manual selection, trace-wise correction factors are extracted. d) For the dynamic traces selected for further analysis, the dwell-time distributions are determined using a Hidden Markov Model approach. e) From the HMM analysis, transition density plots are extracted from the smFRET data.

Next, the recorded traces were analyzed (Supplementary Figure SN5.1b-e) either manually (Section 5.4–5.9) or assisted by neural networks (cf. Supplementary Note 1). Manual evaluation of single-molecule or multi-color FRET traces involves:

(4) the pre-sorting of traces suitable for 1c, 2c-, and 3c-smFRET analyses
(5) determining consecutive regions of the trace for evaluation and for determination of local and global correction factors
(6) Hidden Markov Modeling of smFRET traces to identify underlying states and dwell-times
(7) kinetic evaluation of transition rates and states using transition density plots (TDP), state-wise histograms and dwell-time analyses.

#### 5.2. Camera mapping for FRET traces

In order to extract the fluorescent intensity traces of individual, fluorescently labeled DNA origami structures detected in various channels, an accurate localization and mapping of the detected emission channels across the three cameras or detection areas needs to be achieved. To compensate for potential chromatic aberrations and non-ideal alignment, an image transformation was used to map the corresponding pixels between different cameras/regions onto each other. The associated transformation matrix describing the potential shifts, tilts, etc. was obtained by imaging a calibration pattern on all three detection channels (Supplementary Figure SN5.1a). As a calibration pattern, we typically use a zero-mode waveguide array.

#### 5.3. Trace extraction and background subtraction

After mapping the different detection channels, the location of the individual single emitters needs to be determined and the intensity extracted. When msALEX excitation is used, the alternating laser excitation scheme needs to be taken into account and the intensity traces separated based on both the detection and excitation channels. The most blue-shifted detection channel serves as the reference channel (Channel 1). This refers to the blue excitation, blue channel (BB) for BY-, BR- and BYR-labeled samples or the yellow detection channel with yellow excitation (YY) for YR-labeled samples (Supplementary Figure SN5.1a).

Individual molecules in the reference channel are identified by searching for the brightest spot in the summed projection of the movie. After calculating the central position of the molecule using a wavelet approach^11^, the corresponding position in all other channels is calculated using the transformation matrix. Molecules in the projection images exhibiting detectable intensity in all desired channels are then selected. The intensities and background are extracted using different masks. For the signal, the pixels within an approximate circle of roughly 3 pixels radius around the central coordinates of the molecule are summed together. With a pixel size of 124 nm, the fluorescence signal of a single molecule is accumulated within an area of 614 × 614 nm^2^. For the background, a mask representing roughly a circle with radius of 7.5 pixels (850 nm) and width of 2 pixel centered on the molecule is used. The background is calculated as the median value of all pixels inside the ring-shaped mask and averaged over a five-frame sliding window depending on the excitation cycle and the detection channel. Afterwards, the determined background is scaled to the signal mask and subtracted from the framewise intensity per each channel for each molecule. When analyzing single molecule traces from hand, trajectories which contain molecules within the background mask are discarded. In DeepLASI, these traces are typically discarded in the ‘artifact’ category during the first characterization step.

#### 5.4. Manual trace selection and analysis

The background-corrected fluorescence intensity traces of the individual molecules are then inspected and sorted (Supplementary Figure SN5.1b). The properties of the extracted traces are generally very heterogeneous. This stems from different sources including photochemistry, dye blinking, aggregates and impurities within the sample of different brightness. In all cases, molecules were rejected automatically if they exhibited (1) a low SNR or (2) a brightness that is significantly higher than expected for a single fluorophore (aggregates or impurities). We further classified traces according to

(1) their static and dynamic behavior
(2) the existence of photobleaching steps in the different intensity channels
(3) the order of bleaching steps between the different intensity channels
(4) the degree of labeling efficiency

With the presorted trajectories at hand, we next prepared the data either for (1) correction factor determination to obtain accurate FRET efficiencies (Supplementary Note 5.5) or (2) directly to kinetic and state evaluation based on background-corrected trajectories (1-color data) or apparent FRET efficiencies (2/3-color data; Supplementary Note 5.6). In the first case, we first derived the correction factors per trace and marked regions for trace evaluation by HMM afterwards. In the second case, we manually marked the regions in traces to be analyzed and added them to the ‘HMM’ category.

#### 5.5. Accurate FRET determination

In real smFRET experiments, the intensity of the acceptor signal needs to be additionally corrected for direct excitation of the acceptor fluorophore and spectral crosstalk from donor into the acceptor channel. In addition, the one needs to correct for the difference in the detection sensitivity between the donor and acceptor fluorophores. The correction factors are denoted as:

*de_XY_* for direct excitation of the acceptor fluorophore *Y* during excitation with *X,*
*ct_XY_* for spectral crosstalk from the fluorophore *X* in the detector channel *Y*,
and *γ*_XY_ compensates for differences in detection sensitivities between channels.

We denote the background-corrected intensities as *I_XY_* and the corrected Intensity as *I_XY,corr_*, where *x* stands for the excitation source and *y* for the emission channel, i.e., *I_BR,corr_* denotes the background corrected emission of the acceptor within the red channel (R) after donor excitation in the blue channel (B).

##### Trace-wise and global correction factors

Depending on when individual fluorophores photobleach, some of the correction factors can be extracted from the trace itself. However, in the vast majority of the traces, one cannot extract all correction factors individually. When a trace-wise correction factor is unavailable or unreasonable, the *median* value of the corresponding distribution of trace-wise correction factors for the particular correction factor is used to calculate the accurate FRET values, i.e. a global correction factor. Using traces that were presorted and categorized as ‘Blue dye bleached’ (or ‘yellow / red dye bleached’, respectively), we first determined the trace-wise correction factors for direction excitation *de_XY_* and spectral crosstalk *ct_XY_*. Having corrected the background-corrected intensities against both contributions, we next determined the trace-wise correction factor *γ_XY_*.

To derive the contribution of spectral crosstalk from the donor channel *X* in the acceptor channel *Y*, we determine the trace-wise correction factor *ct_XY_* using the intensity information after photobleaching of the acceptor:

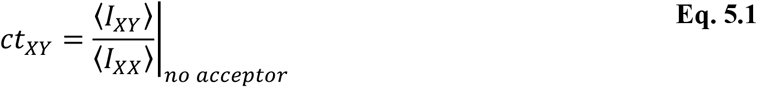

Here, 〈*I_XX_*〉 refers the mean donor intensity and 〈*I_XY_*〉 to the mean acceptor intensity after donor excitation in the region of the trace where there is no acceptor fluorescence.

Similarly, we determined the correction factors for direct excitation of the acceptor during donor excitation using traces in which the donor fluorophore bleached first:

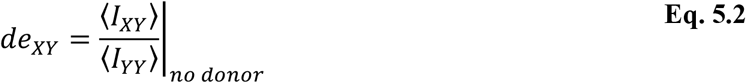

where 〈*I_XY_*〉 and 〈*I_YY_*〉 describes the mean acceptor emission after donor excitation or acceptor excitation, respectively.

Lastly, we determined the detection correction factors *γ_XY_* compensating for differences in detection sensitivities between different channels. For this, we used traces where the acceptor photobleaches before the donor. The acceptor intensity is first corrected for direct excitation *de_XY_* and spectral crosstalk *ct_XY_*. We then derive the detection correction factor *γ_XY_* per trace from the ratio of changes in donor and acceptor emission before and after photobleaching of the acceptor. The correction factors are denoted as:

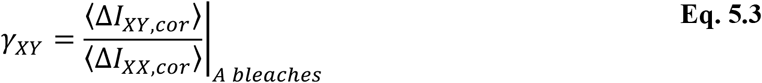

where 〈Δ*I_XX,cor_*〉 and 〈Δ*I_XY,cor_*〉 refer to the intensity difference for the mean donor and acceptor emission after donor excitation before and after acceptor photobleaching.

##### Data Correction

Once all correction factors are determined, every trace is corrected using the local, trace-wise correction factors, when available and suitable. Otherwise, the global correction factor is used. In three-color experiments, the corrected FRET efficiency for *E_YR_* is calculated first since it is required for subsequent corrections. Upon yellow excitation, the same approach is used as for two-color FRET experiments:

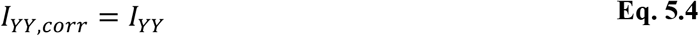

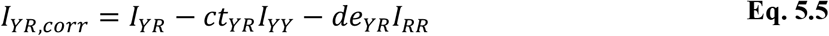

The corrected FRET efficiency is then given by the ratio of both corrected intensities

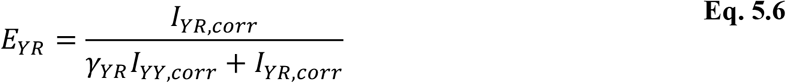

For the BY FRET pair, the fully corrected intensities after blue excitation read as:

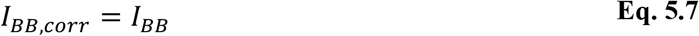

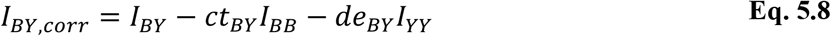

The accurate BY FRET efficiency follows equation 5.5 with an additional term which takes into account the reduction in brightness of the yellow dye due to the FRET process between the YR pair:

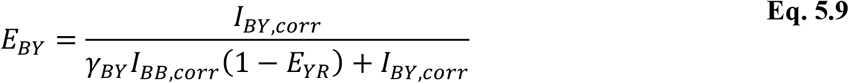

The intensity of the red fluorophore after blue excitation needs to be corrected against direct excitation, contributions of both the blue and yellow dye due to crosstalk into the red channel and due to cascading of FRET from the blue dye over the yellow dye into the red channel:

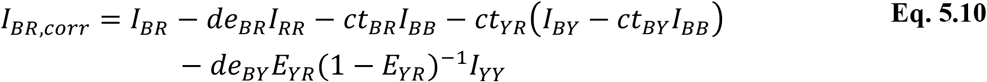

The accurate FRET efficiency of the BR FRET pair is then given by:

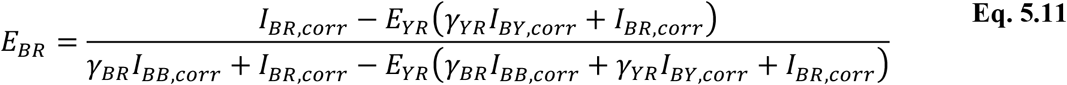

#### 5.6. Hidden-Markov modeling

The kinetics and underlying states within the selected trajectories, i.e., either smFRET or intensity traces, were evaluated using Hidden Markov Modeling. The input data of both assays vary between 0 and 1. We anticipate that every molecule undergoes transitions between a fixed numbers of conformations described by a discrete number of states *q_i_* (*i* = 1, … *Q*). The probability to observe the molecule while remaining in the original state or transiting into either of the other states is given by an exponential decay independent on the dwell time. For a system with Q states in total, the transition probability matrix 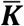 comprises *Q* × (*Q* − 1) independent transition probabilities *k_ij_* describing the likelihood for going from state *i* to state *j*. Here, it is a prerequisite for the Markovian process, that the row-wise sum of transition probabilities is normalized to 1. For a Hidden-Markovian process, the state sequence is not directly observable but buried in random noise of the system. It can only be inferred from measured observables ***x**_t_*, i.e., the single-molecule trajectory, with a length of *T* data points. Here, the emission probabilities 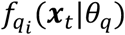 serve as parameter to represent the relative likelihood for observing a specific FRET value (or intensity value) for a given set of model parameters *θ_q_* and the molecule being in state *q_i_*. We model the emission probability as a Gaussian distribution to describe the probability density function of a state *q_i_*.^12^

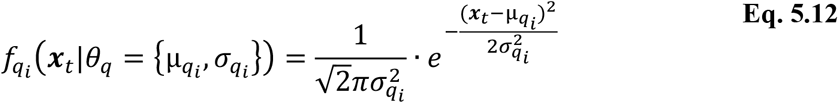

The parameters estimators are: the mean value 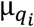 and covariance 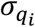

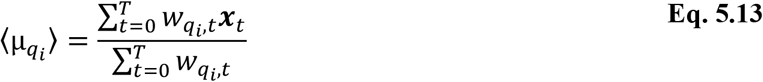

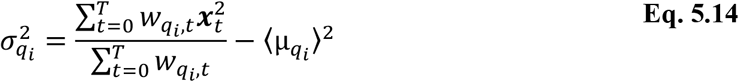

For this we introduced the relative occurrence probability 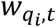, i.e., the conditional probability 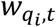 of being in state *q_i_* given the data ***x**_t_* at a time t, which is linked to the fraction of time *W_q_*

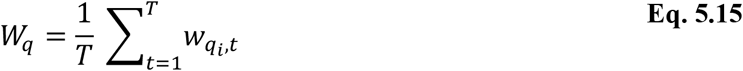

and emission probability emission probability 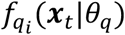

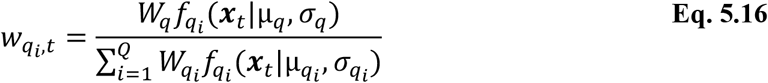

To describe the underlying Markov chain in individual time trajectories, we describe the system in terms of emission and transition probabilities. The likelihood for switching between states *q_i_* given the observed FRET data ***x**_t_*, is simply obtained by multiplying the probability for individual time points *t* and is given as^13^

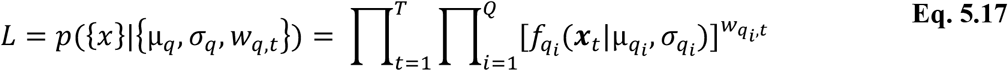

To derive the transition density matrix, the HMM needs to be trained on the experimental single-molecule data. This working step can be carried out in two different approaches: the analysis can be carried out either (1) trace-wise or (2) globally. In the first case, the transition probability and emission probability are optimized for each individual trace while, in the second case, one uses a shared single transition probability matrix and parameters for the emission probability for all trajectories together.

We used a trace-wise approach when analyzing 1-color and apparent FRET traces as the exact value of the state can be shifted due to the above-mentioned background contributions.

#### 5.7. Evaluation of involved FRET states and interconversion rates

The last step involves the visualization of determined rates, i.e., dwell times, and states determined from the FRET and / or normalized intensity traces. We employed so called transition density plots (TDPs), which depict each transition that was identified by the HMM algorithm or the Deep-LASI state classifier in the recorded time traces as a single event in a 2D diagram. The diagram, hence, depicts and links the FRET value before and after an identified transition visually. In the case of 1-color data, we normalized the traces between the minimal and maximal value of observed counts of all measured single traces. The TDPs were generated as described by McKinney et al., i.e. all transitions are depicted as summed up two-dimensional Gaussian functions with an amplitude equal to the total number of transitions and a fixed variance of 0.0005.^12^

### SUPPLEMENTARY NOTE 6: DETAILS OF DEEP-LASI ANALYSES

#### 6.1 Results for the three-color, two-state DNA origami structure with different binding site lengths

The three-color DNA origami structures were measured with four different lengths of complementary DNA for the two binding sites. The two binding sites contained the identical DNA sequence and lengths. The dwell time distributions determined from the state classifiers of Deep-LASI for the different three-color DNA origami structures are shown in Supplementary Figure SN6.1. The same analysis workflow was followed for each sample: a fully automated categorization and prediction of state occupancy in traces labeled as ‘dynamic’ were performed with Deep-LASI followed by a manual selection of the different states and fit to a mono-exponential function. These experiments confirm that Deep-LASI is capable of extracting mono-exponentially distributed dwell times over a large range of kinetic rates.

**Figure SN6.1:**
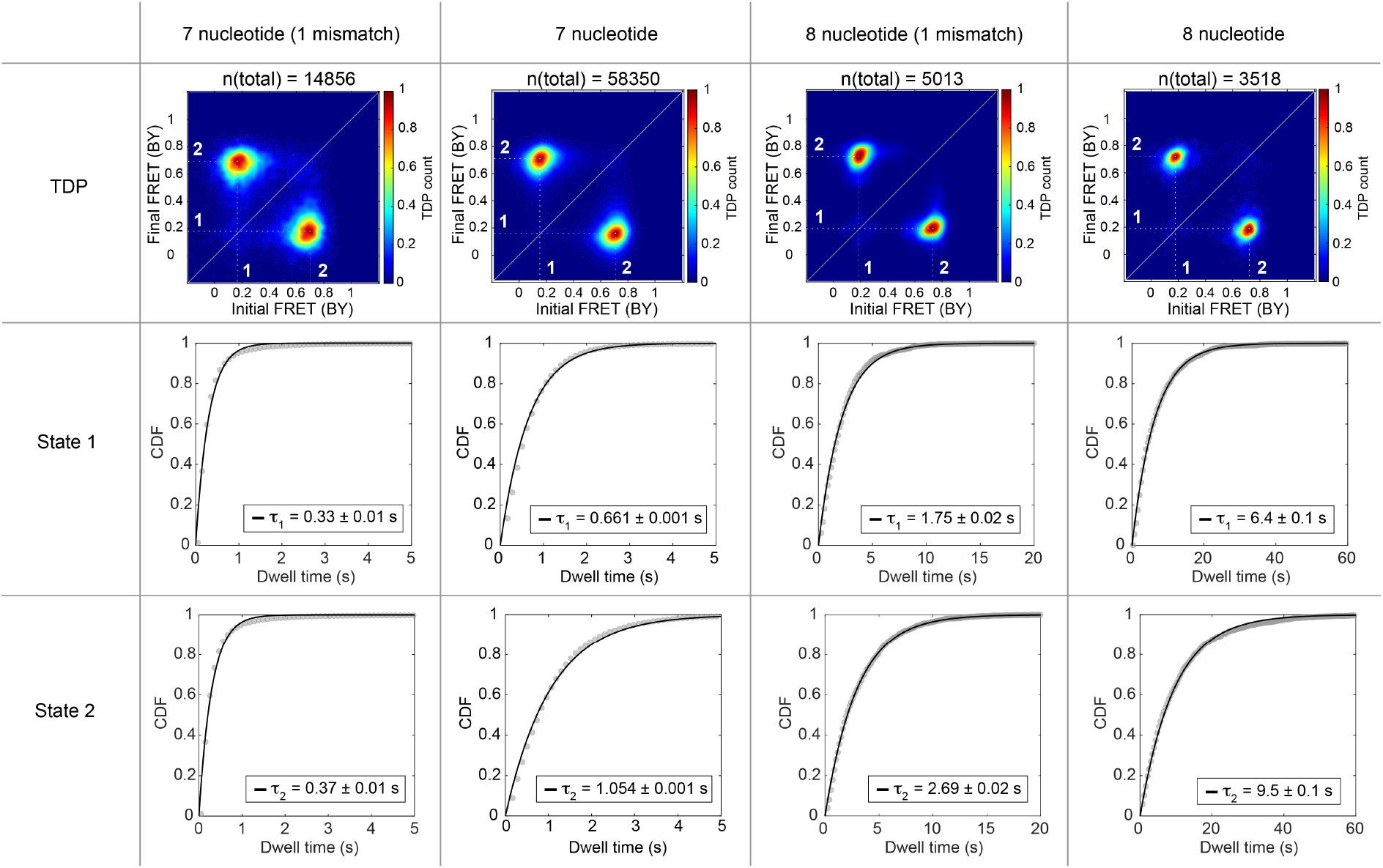
Dwell-time distributions of the three-color, two-state DNA origamis with different binding site lengths. Each row corresponds to a specific state and each column depicts the TDPs (top) and dwell-time distributions (middle, bottem) extracted from the uncorrected blue-yellow transition density plots and fitted with a mono-exponential for each binding site length.

#### 6.2 Kinetics of the three-color, three-state DNA origami

From the three-color, three-state DNA origami with 7 nt binding strands at positions 6 and 12 o’clock and a 7.5 nt complementary binding strand at 9 o’clock. Three populations were extracted automatically from the traces identified by Deep-LASI as dynamic. The dwell-time distributions of all 6 populations observed in the blue/yellow TDP plot (Figure 5c) were extracted manually and fit with an exponential function (Supplementary Figure SN6.2). The dwell times of each state are in excellent agreement with the two-color, three-state DNA origami sample (Supplementary Figure SN6.3), indicating that the additional blue dye in close proximity of state 2 does not influence the kinetic rates.

**Figure SN6.2:**
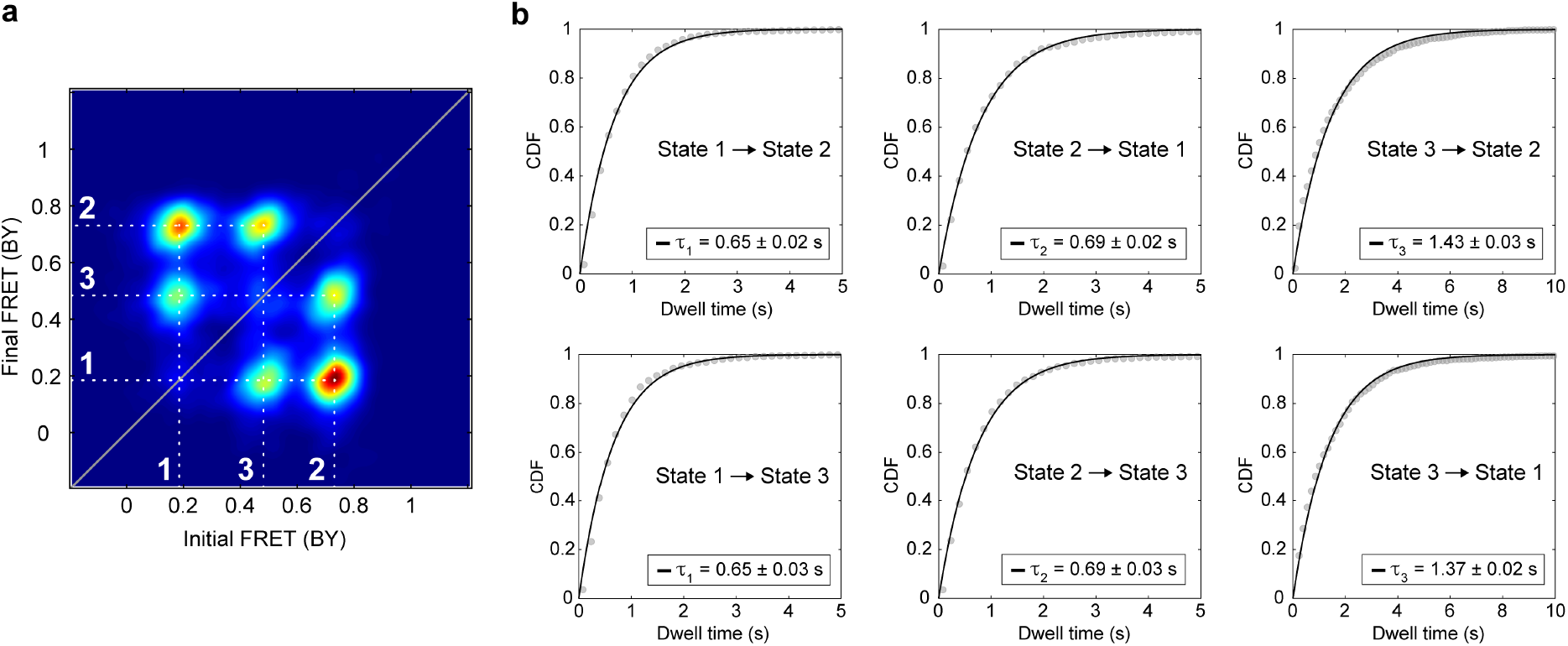
Dwell-time distributions of the 3-color 3-state DNA origami. **(a)** The blue/yellow transition density plot and **(b)** the dwell-time distributions extracted from the BY-TDP and fit using a mono-exponential.

#### 6.3 Results for the two-color, three-state DNA origami structure

Next, we tested the performance of Deep-LASI on a more complex, two-color, multi-state system by introducing a third binding site on the DNA origami (Supplementary Figure SN6.3a) and increasing the average transition rates. In contrast to the two-state system described above, State 1 and State 2 at the 6 o’clock and 12 o’clock positions are now characterized by 7 nt binding sites in the three-state DNA origami. The added State 3 at 9 o’clock has a 7.5 nt overhang. In the example trace shown in Supplementary Figure SN6.3b, Deep-LASI extracts the dynamic section and identifies all transitions between the three states summarized in the TDP of apparent FRET efficiencies (Supplementary Figure SN6.3c). As expected, the FRET efficiency of state 1 (0.83) and state 2 (0.21) do not change significantly compared to the two-state system. In addition, a third state with an apparent FRET efficiency of 0.31. However, as states 2 and 3 show a similar distance to the acceptor, the states and thereby the transitions are not easily separable. When looking at the dwell-time distributions, the transition out of state 1 is not affected by the degeneracy of states 2 and 3. However, the transition rates from state 2 or 3 to state 1 differ significantly due to the different binding site lengths and can only be extracted using a bi-exponential fit (Supplementary Figure SN6.3d). From the TDP, we can also extract the transitions between states 2 and 3. The transition from state 2 to state 3 can be well described by a monoexponential distribution whereas the reverse transition from state 3 to state 2 has a second component due to the difficulties of clearly separating the different states.

From the single molecule trajectories, Deep-LASI also extracts the regions of the trace that can be used for determining the different correction factors. The FRET correction factor distributions determined by Deep-LASI are shown in Supplementary Figure SN6.3e and are consistent with the correction factors of the two-state DNA origami data set shown in Figure 3f. The framewise apparent smFRET histogram is shown in Supplementary Figure SN6.3f (top, gray). In this histogram, states 2 and 3 merge into one degenerate state (0.27) due to heterogeneous broadening of the two populations. After correction (Supplementary Figure SN6.3f, top, orange), the degeneracy is decreased and the low-FRET peak broadens. However, they are still not clearly separable. It is only after using the state-label information, which allows us to average the state FRET efficiencies that the two low-FRET populations become distinguishable and the individual FRET populations observed (Supplementary Figure SN6.3f).

**Figure SN6.3:**
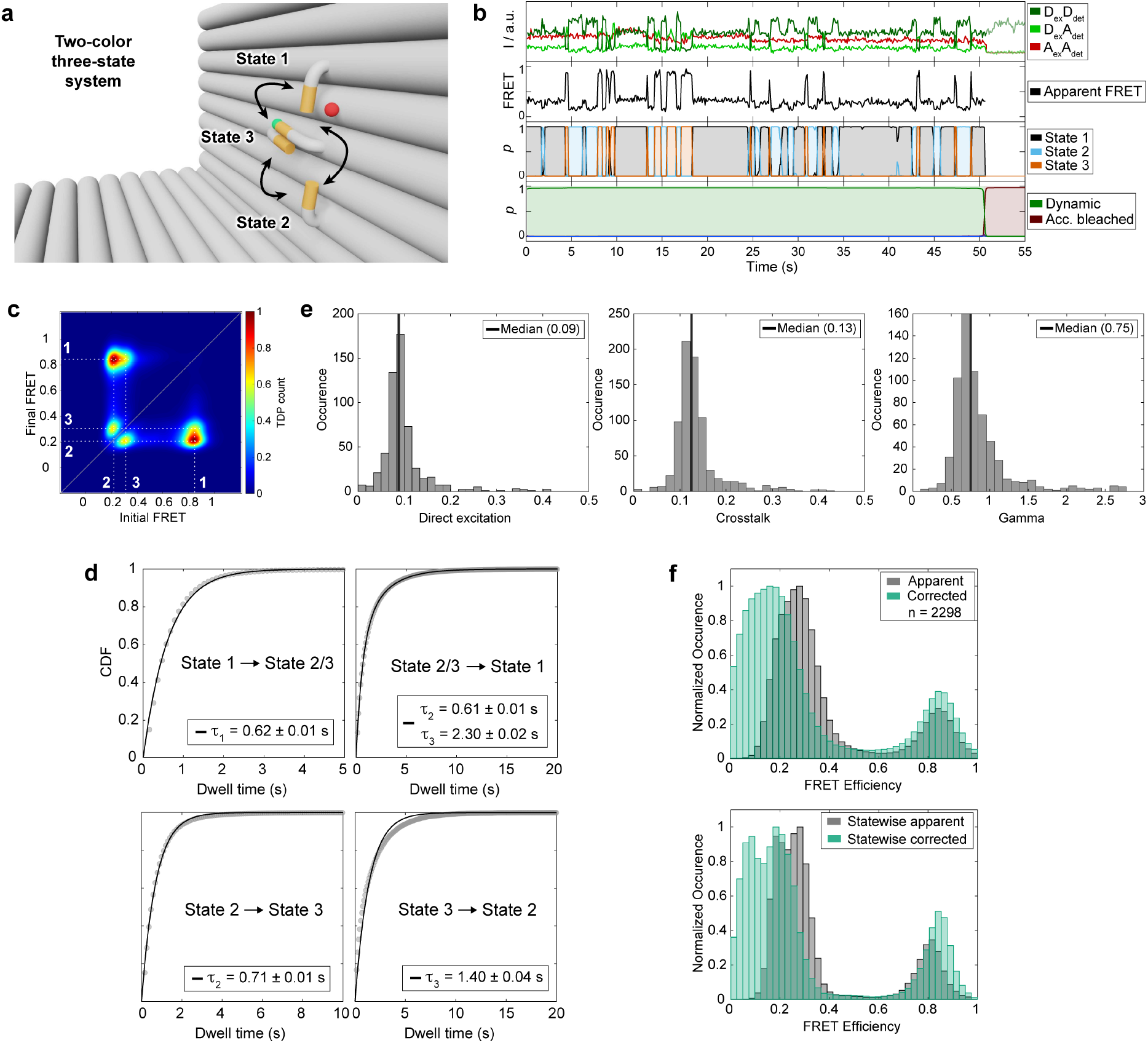
Analysis of 2-color, 3-state DNA origami measurements. **(a)** Zoom-in of the L-shaped DNA origami structure with three binding sites. FRET is expected between a high FRET state 1 (12 o’clock), a low FRET state 2 (6 o’clock), and an intermediate FRET state 3 (9 o’clock). **(b)** A representative single molecule intensity trace and FRET trajectory. The upper panel shows the intensity in the yellow and red channels after yellow excitation and the red intensity after red excitation. The middle panel shows the corresponding FRET efficiencies for the dye pair. The third and fourth panels show the output of the Deep-LASI analysis for state-transition and trace classification respectively. **(c)** The TDP of the apparent FRET efficiency states are shown. Interconversion between three conformations with apparent FRET efficiencies of 0.21, 0.31 and 0.83 are observed. The three states are labeled in white. **(d)** Exponential fits of the dwell time distributions for all states are plotted. The transitions from state 2 and 3 to state 1 were pooled together due to the high overlap and fit with a bi-exponential function. While the dwell time of state 2 in the bi-exponential fit is close to the dwell time extracted from the single population (state 2 to state 3), the dwell time of state 3 is significantly overestimated compared to the single population of transitions from state 3 to state 2. **(e)** Correction factors for direct excitation, crosstalk and gamma extracted by Deep-LASI. **(f)** *top* Frame-wise weighted state-wise smFRET histograms of apparent and accurate smFRET efficiencies. A broadening of the low-FRET population is observed as the correction of the FRET efficiency begins to lift the degeneracy. *bottom* Plotting the framewise-weighted statewise smFRET histograms of apparent and accurate FRET efficiencies improves the contrast. Three peaks are now observable with corrected FRET efficiencies of 0.09 and 0.84 (in line with the two-state system), and a new third state at 0.19.

#### 6.4. Analysis of kinetic data from the kinsoft challenge

We tested the performance of Deep-LASI on datasets provided by a recently published multi-laboratory software comparison study for extracting kinetics from smFRET data. As we contributed to this study using conventional HMM, we chose to analyze the datasets that did not require additional human input for interpretation of the data. The results are shown in Supplementary Figure SN6.4. Deep-LASI returned values corresponding to the ground truth for the simulated dataset and close to the average values obtained for the experimental dataset.

**Figure SN6.4:**
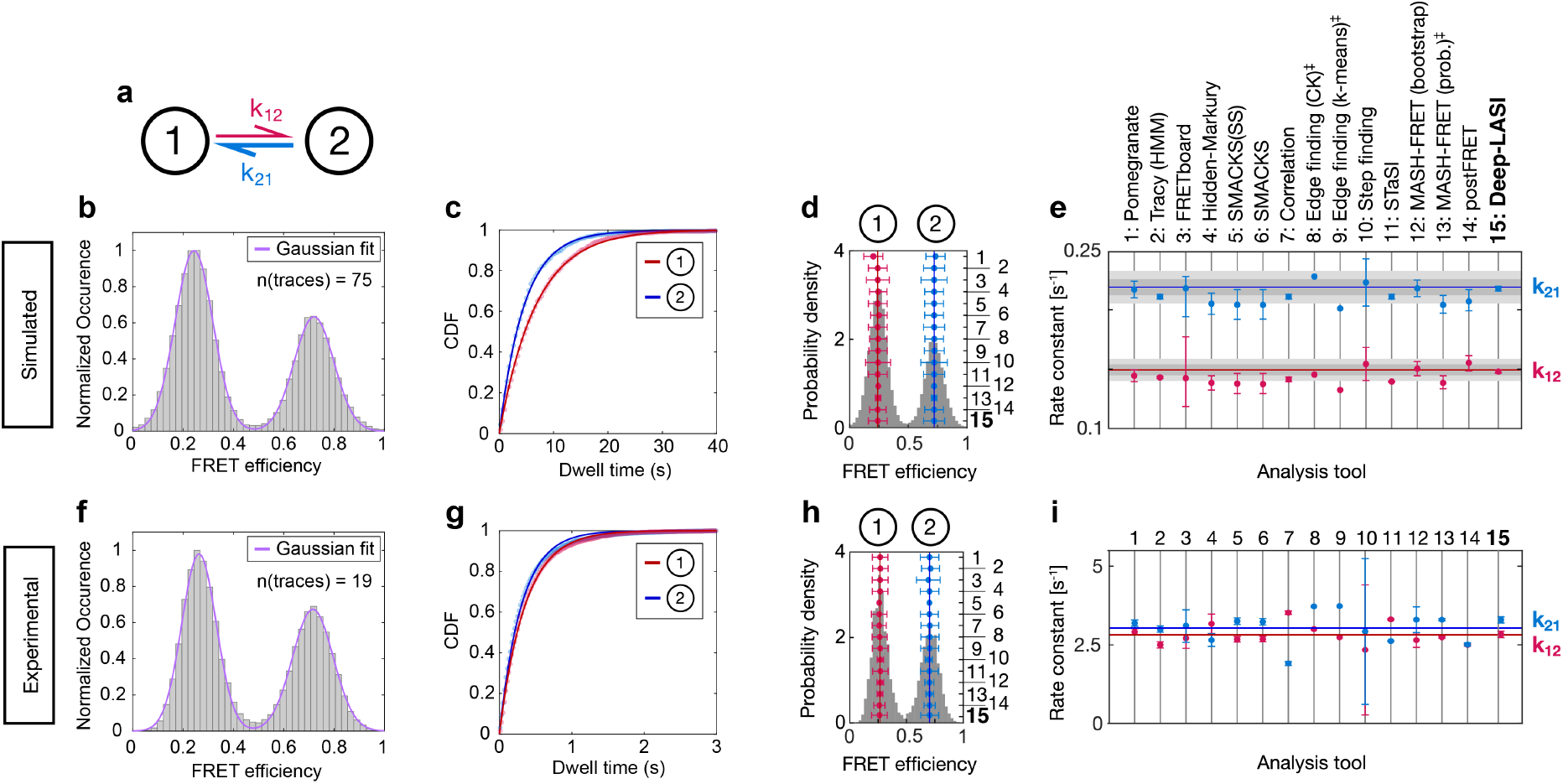
Kinetic analysis of datasets from the kinetic software challenge. **(a)** An illustration of the kinetic two-state model connected by forward and backward rate constants: k_12_ and k_21_. **(b)** A framewise FRET efficiency histogram (gray) of the simulated data extracted by the trace classifier. A Gaussian fit to the two populations are shown in magenta. **(c)** A mono-exponential dwell time distributions of the data in (b) obtained from the state-transition classifier. **(d)** The ground truth FRET histogram (gray) with state assignments labeled at the top and the inferred average FRET efficiencies in red and blue. Numbers on the right axis refer to the analysis tools specified in (e). Vertical lines indicate the mean over all tools. The error bars represent the standard deviations returned from the different analysis routines. **(e)** Rate constants and uncertainties inferred from the dataset in (d) by different labs using the respective analysis tools. The ground truth (GT) is indicated by the horizontal red and blue lines, the intrinsic uncertainty of the dataset is represented by dark gray (1σ) and light gray (2σ) intervals. **(f)** A framewise smFRET efficiency histogram (gray) of the experimental data extracted by the trace classifier. **(g)** The dwell-time distributions and corresponding mono-exponential fits of the data in (f) obtained from the state-transition classifier. A Gaussian fit to the two populations are shown in magenta. **(h)** A smFRET histogram of preselected traces from panel (h) where photobleaching and photoblinking contributions have been removed. State 1 is labeled in red and state 2 in blue. The vertical lines indicate the average value returned from analysis routines 1-14. The legend for the analysis routines is given in (e). The error bars represent the standard deviations returned from the different analysis routines. **(i)** Inferred rate constants from the experimental dataset in (h). The respective analysis tools are specified in (e). Horizontal red and blue lines indicate the mean of the inferred kinetic rate constants from analysis tools 1-14. The legend for the analysis routines is given in (e)

### SUPPLEMENTARY NOTE 7: DNA SEQUENCES

The details of the L-shaped DNA origami structures are described here in detail. The structures were previously published by Tinnefeld et al.^14,15^. As a scaffold, we used the p8064 scaffold derived from M13mp18 bacteriophages. An overview of all designed DNA origami structures including name, the strand IDs of the introduced modified staple strands as well as the binding sites is given in Supplementary Table S7.1. The unlabeled staple strands are specified in Supplementary Table S7.2, staple strands with biotin modifications for surface immobilization are listed in Supplementary Table S7.3 and staple strands with fluorescent modifications for single-molecule FRET are summarized in Supplementary Table S7.3.

The L-shaped DNA origami structures are made of 252 ssDNA staple strands annealed to a circular complementary ssDNA scaffold strand of 8064 nucleotides. The three fluorophores ATTO488, Cy3b and ATTO647N are introduced into the structures, by replacing the unlabeled ssDNA strands L7, L8 and L9 (Supplementary Figure SN7.1) with strands containing the appropriate label (Supplementary Table SN7.4). Binding sites for the L7-attached tether strands consisting of different lengths are introduced at position L5 and L6 for the 2 state systems with low and high FRET values and different binding rates, even with identical sequences. We refer to the binding site for staple strand L5 as 6 o’clock and staple strand L6 as 12 o’clock. For generating a 3 state FRET system, an additional binding site was introduced on staple strands L14 at 9 o’clock. In addition, for the implementation of the 9 o’clock binding site, the staple strands L12 and L13 are replaced by L12-13-I, L12-13-II and L12-13-III (Supplementary Table SN7.4). All samples share biotinylated attachment sites at positions L1-L4 (Supplementary Table SN7.3).

**Figure SN7.1:**
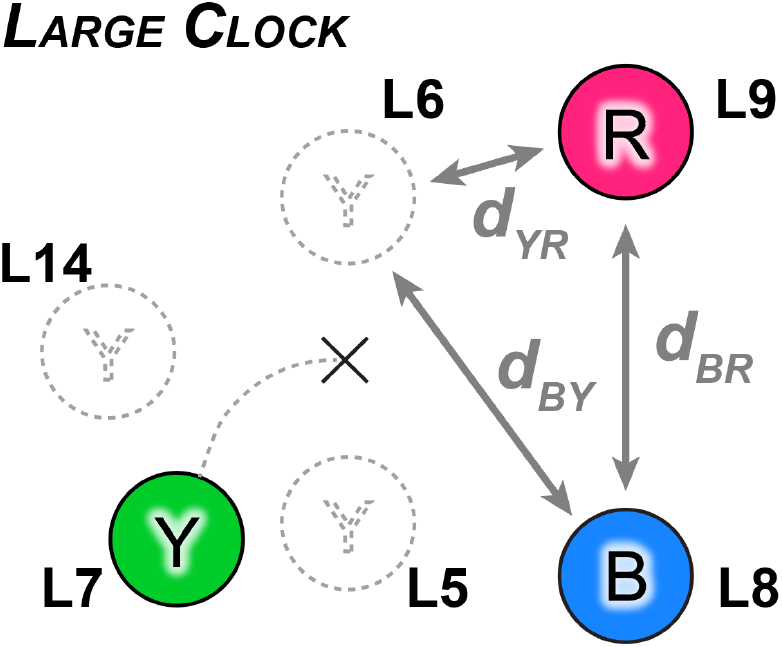
Schematic of the replacement staple strands forming the 3 state, 3color FRET clock on the L-origami. The strands either carry one of the three fluorophores (L7, L8, and L9) or represent a binding site at 6, 9 and 12 o’clock (L5, L14 and L6 respectively).

**Supplementary Table SN7.1.** The applied nomenclature used for the designed L-shaped DNA origami structures with the corresponding staple strand IDs that carry the fluorescent dyes or the attachment of the pointer. The laser excitation scheme for the 3cFRET B-Y-R samples involves excitation at 488, 561 and 640 nm.

| # | Name | Blue | Yellow | Red | Binding sites |  |  | Replaced |
| --- | --- | --- | --- | --- | --- | --- | --- | --- |
| 1 | BYR-Pos6/12-6.5nt | 488-L9 | 561-L7 | 640-L8 | 6.5nt-L5 | 6.5nt-L6 | --- | --- |
| 2 | BYR-Pos6/12-7nt | 488-L9 | 561-L7 | 640-L8 | 7nt-L5 | 7nt-L6 | --- | --- |
| 3 | BYR-Pos6/12-7.5nt | 488-L9 | 561-L7 | 640-L8 | 7.5nt-L5 | 7.5nt-L6 | --- | --- |
| 4 | BYR-Pos6/12-8nt | 488-L9 | 561-L7 | 640-L8 | 8nt-L5 | 8nt-L6 | --- | --- |
| 5 | BY-Pos6/12-7.5nt | 488-L9 | 561-L7 | --- | 7.5nt-L5 | 7.5nt-L6 | --- | --- |
| 6 | BR-Pos6/12-7.5nt | 488-L9 | --- | 640-L8 | 7.5nt-L5 | 7.5nt-L6 | --- | --- |
| 7 | YR-Pos6/12-7.5nt | --- | 561-L7 | 640-L8 | 7.5nt-L5 | 7.5nt-L6 | --- | --- |
| 8 | YR-Pos6/9/12 | --- | 561-L7 | 640-L8 | 7nt-L5 | 7nt-L6 | 7.5nt-L14 | L12-13-I-III |
| 9 | BYR-Pos6/9/12 | 488-L9 | 561-L7 | 640-L8 | 7nt-L5 | 7nt-L6 | 7.5nt-L14 | L12-13-I-III |

**Supplementary Table SN7.2.**
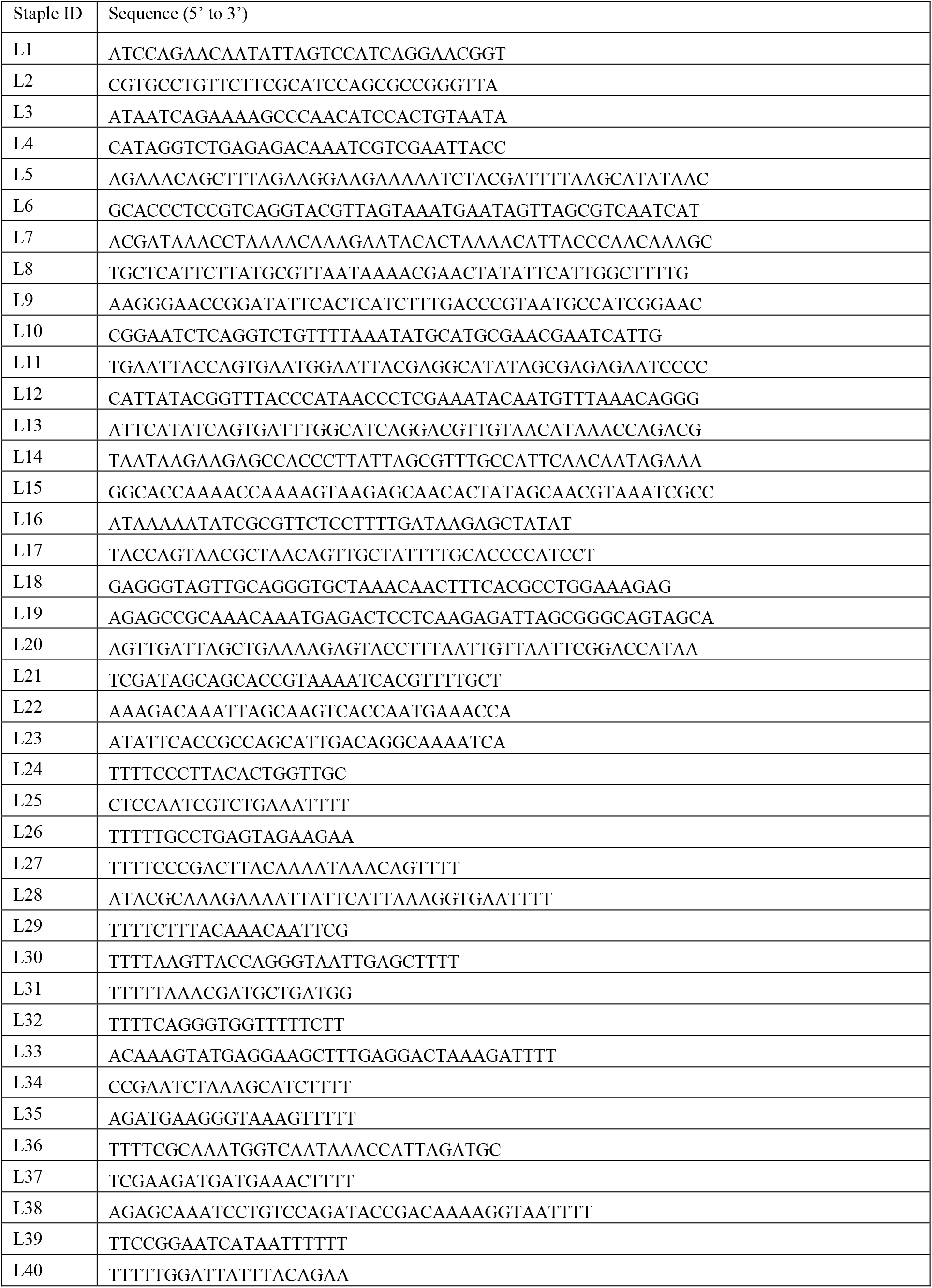
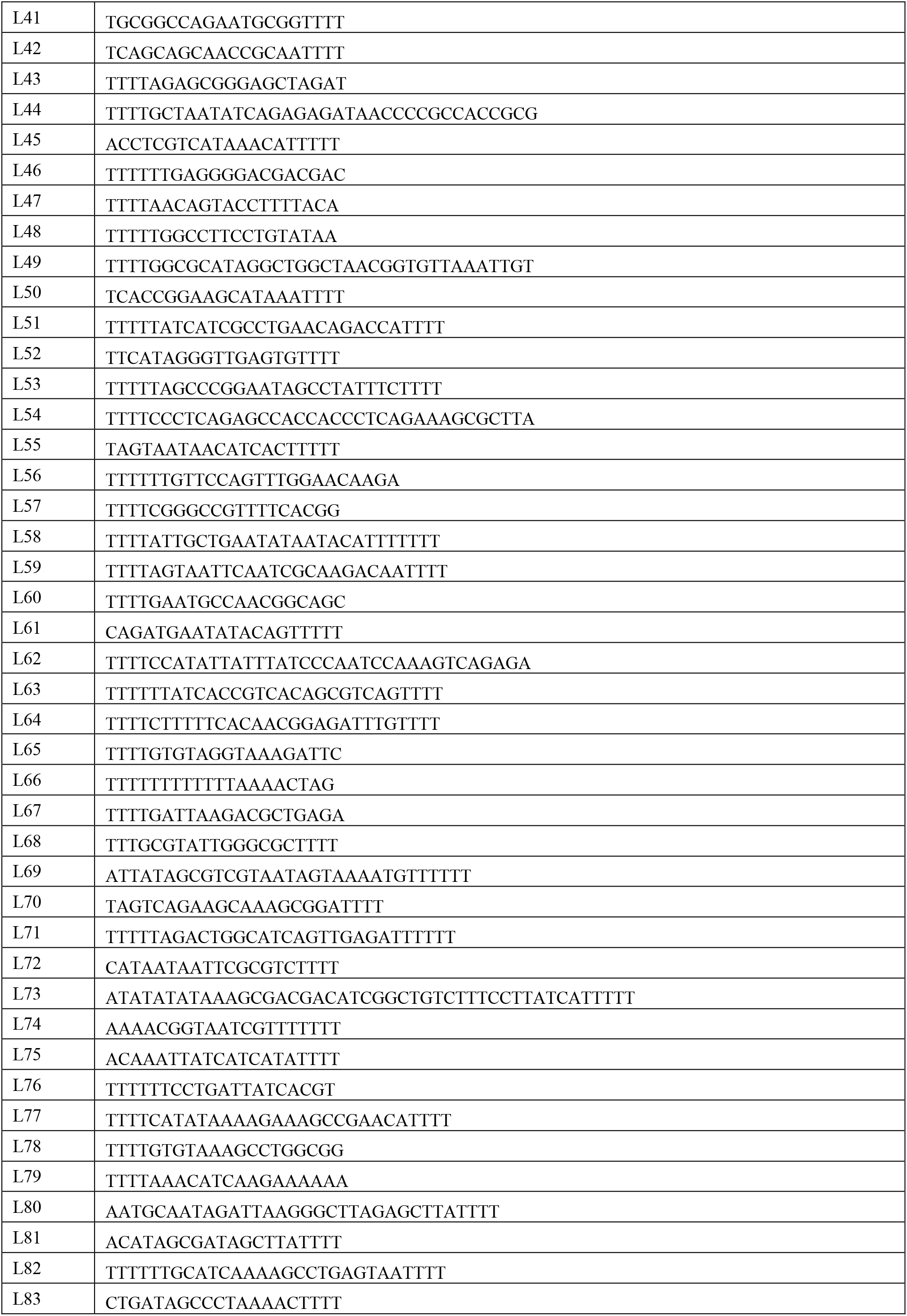
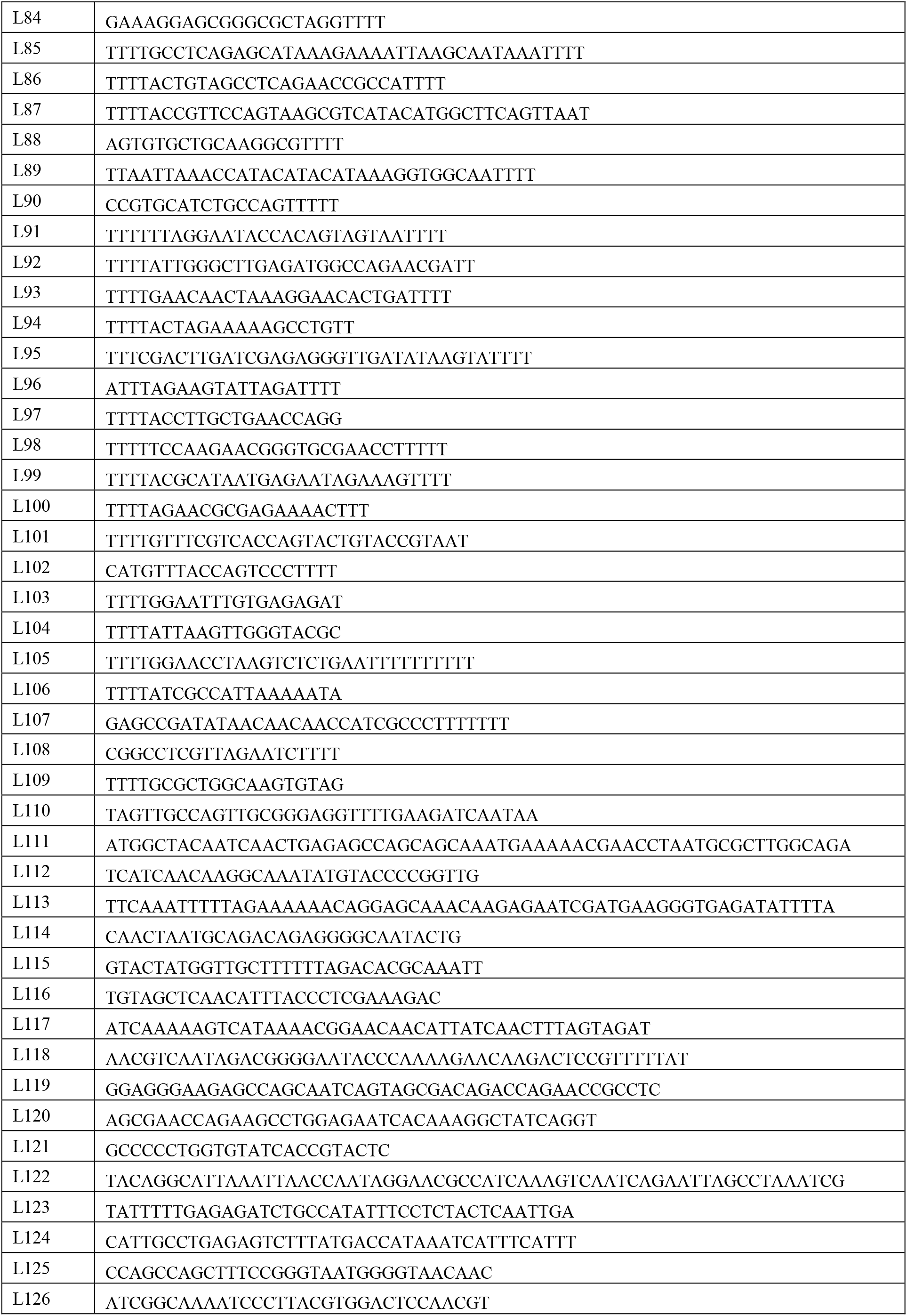
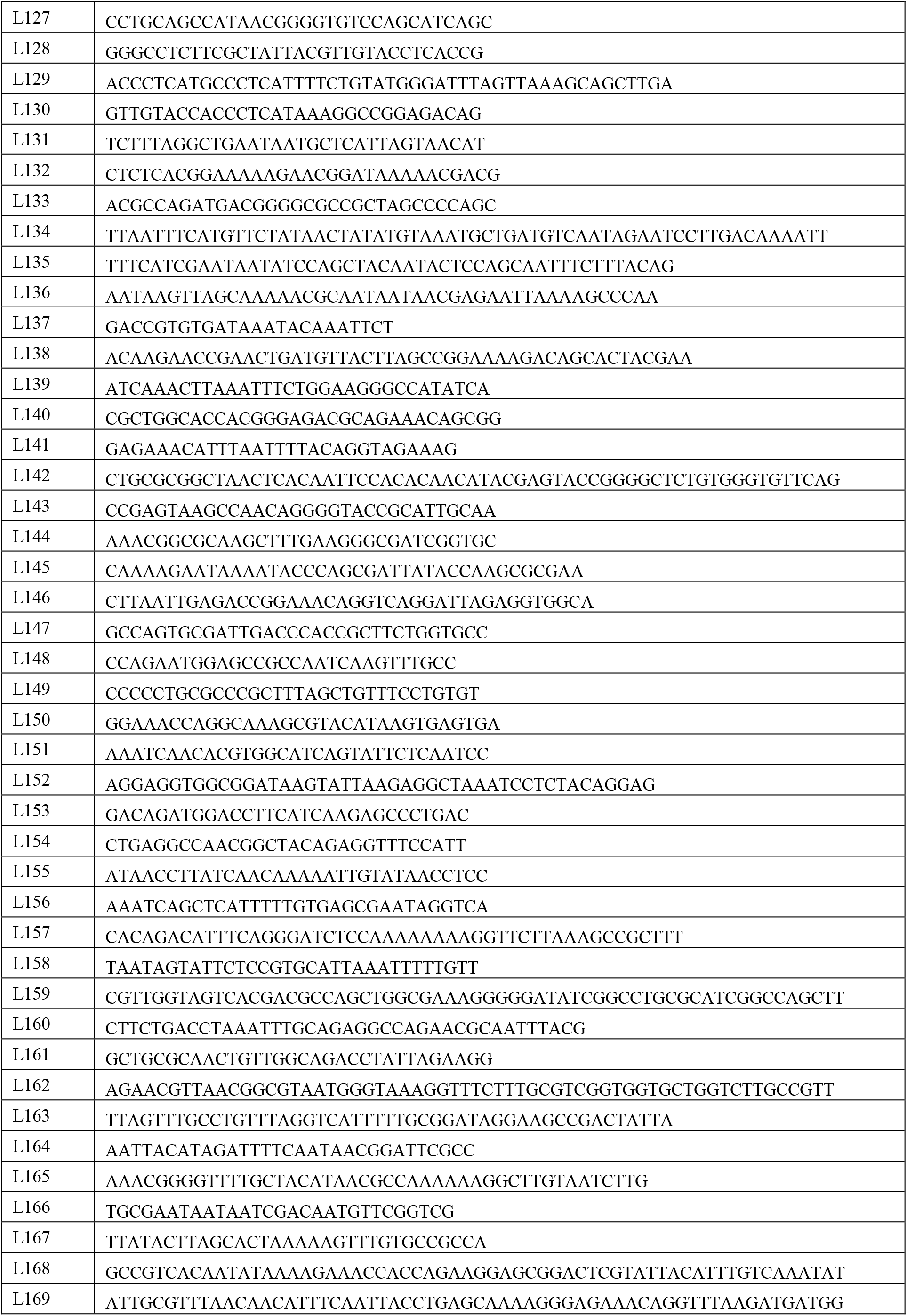
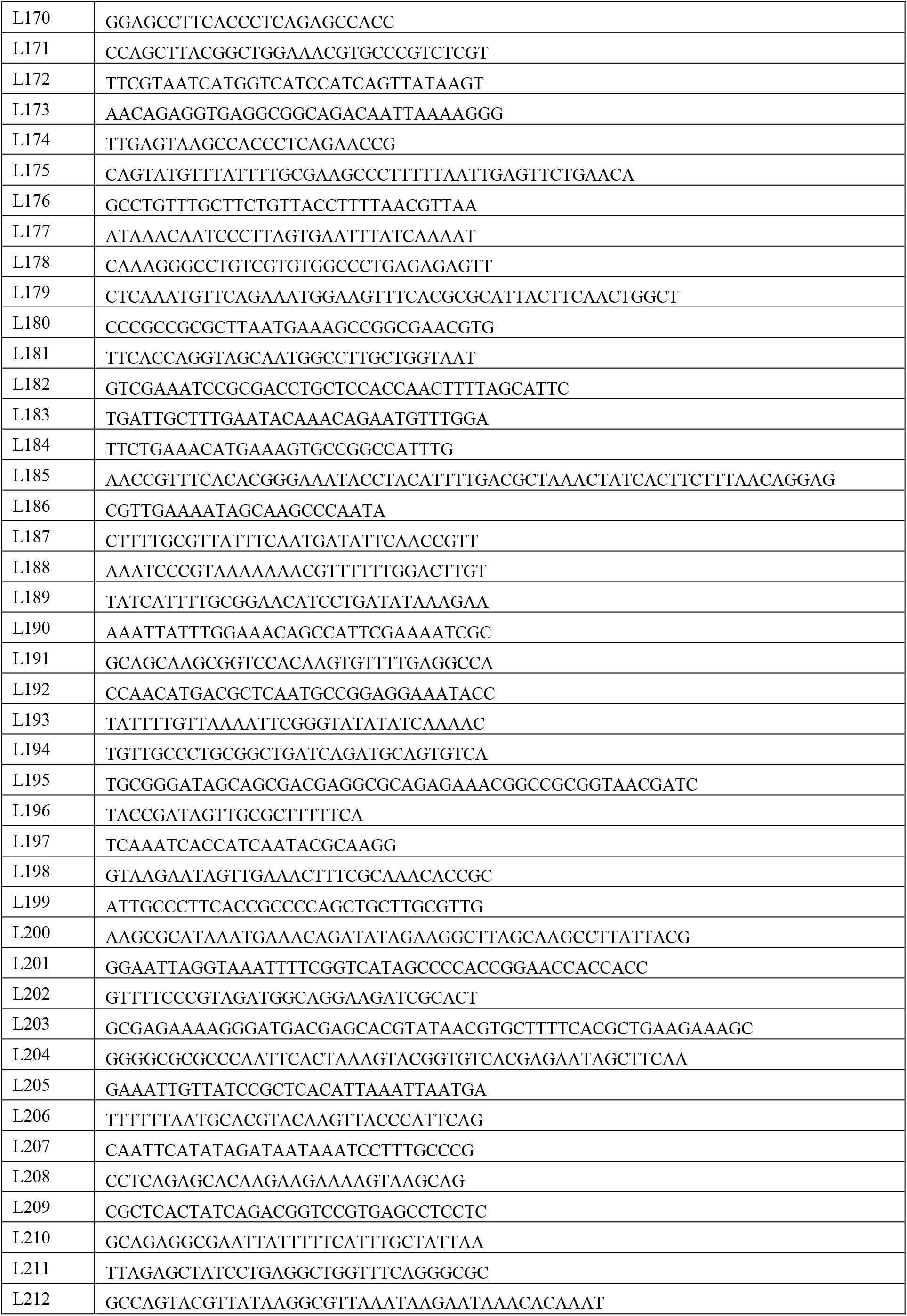
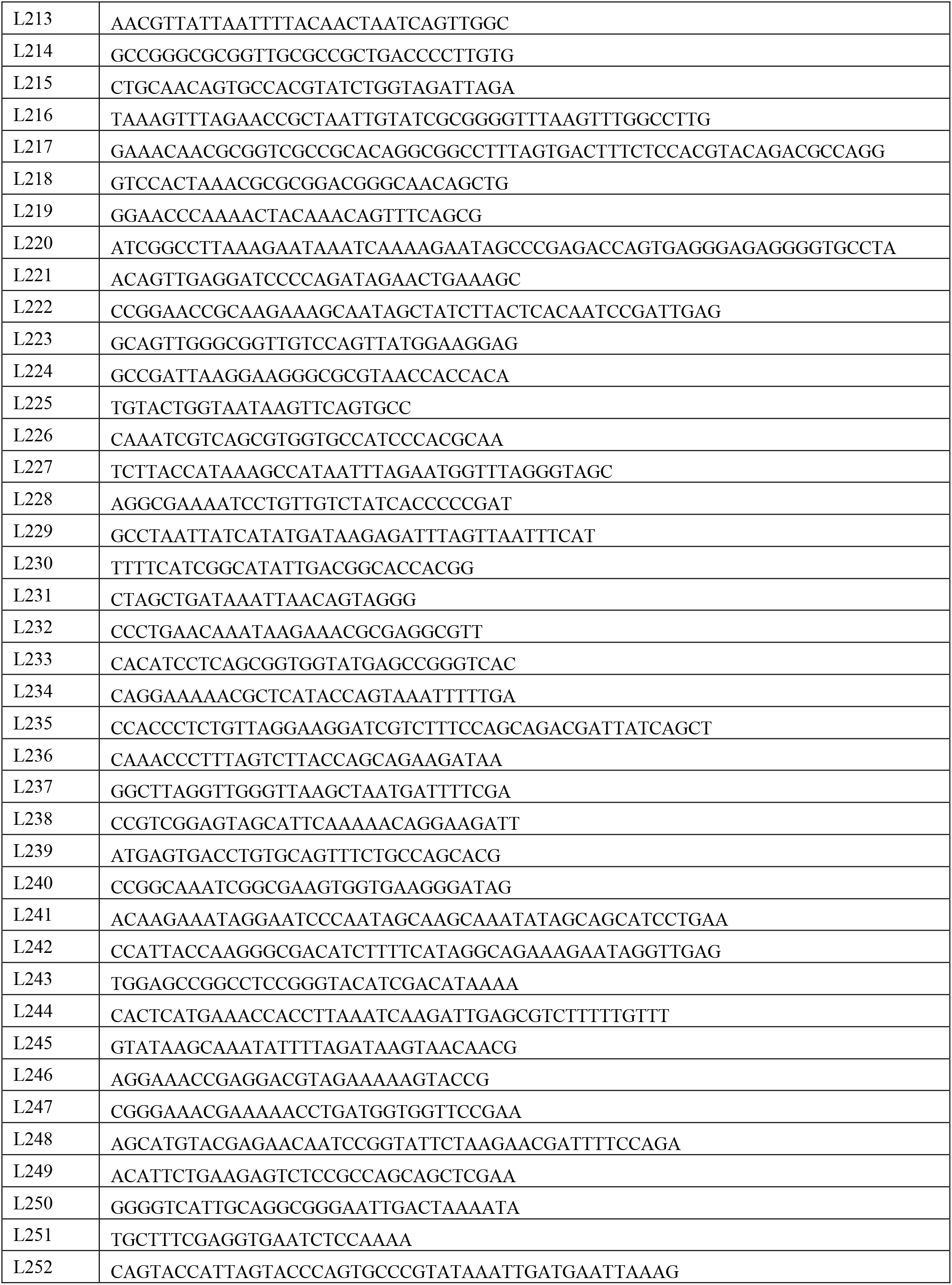
Unmodified staple strands used for the L-shaped DNA origami structure given from the 5’ to 3’ end. All oligonucleotides were purchased from Integrated DNA Technologies.

**Supplementary Table SN7.3.**
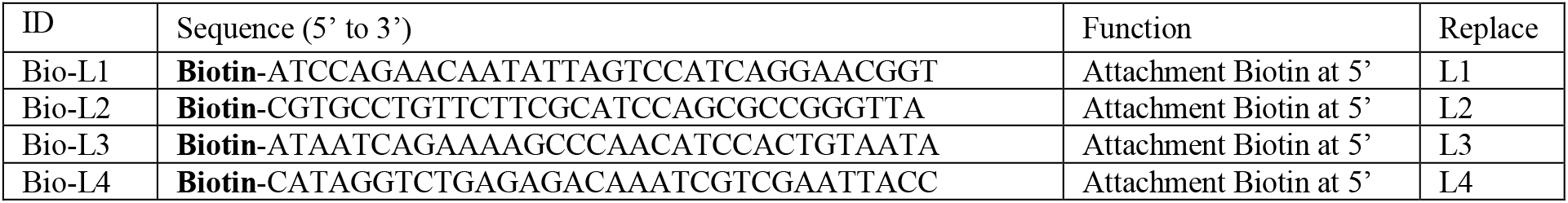
Modified staple strands given from the 5’ to 3’ end for the L-shaped DNA origami structures used. The biotin was used for surface-immobilization via a biotin/avidin interaction. All oligonucleotides were purchased from Biomers.

**Supplementary Table SN7.4.**
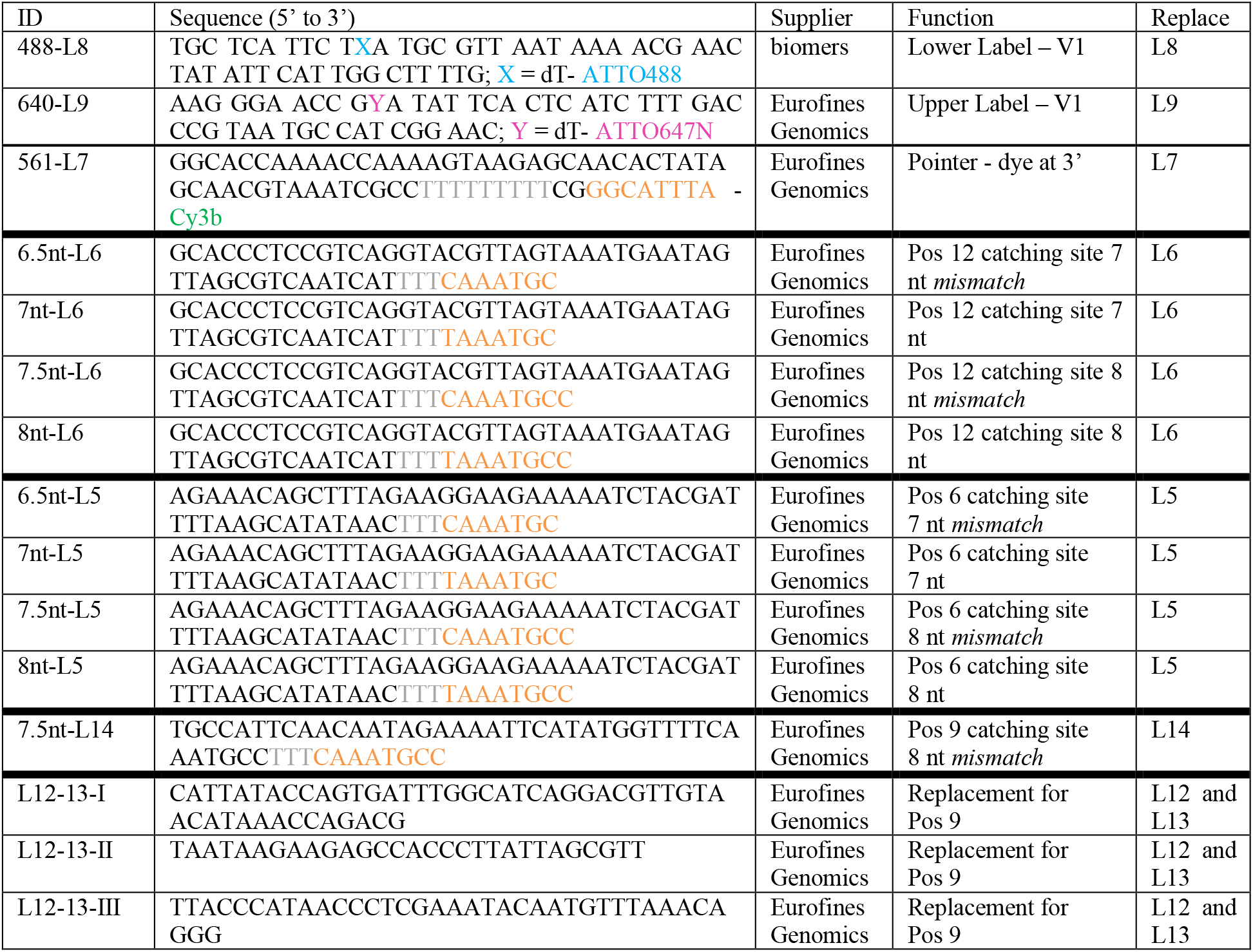
Modified staple strands given from the 5’ to 3’end for the fluorescently-labeled L-shaped DNA origami structures.

